# Ca^2+^-inactivation of the mammalian ryanodine receptor type 1 in a lipidic environment revealed by cryo-EM

**DOI:** 10.1101/2021.11.14.468550

**Authors:** Ashok R. Nayak, Montserrat Samsó

## Abstract

Activation of the intracellular Ca^2+^ channel ryanodine receptor (RyR) triggers a cytosolic Ca^2+^ surge, while elevated cytosolic Ca^2+^ inhibits the channel in a negative feedback mechanism. Cryo-EM carried out under partially inactivating Ca^2+^ conditions revealed two conformations of RyR1, an open state and an inactivated state, resolved at 4.0 and 3.3 Å resolution, respectively. RyR1s were embedded in nanodiscs with two lipids resolved at each inter-subunit crevice. Ca^2+^ binding to the high affinity site engages the central (CD) and C-terminal domains (CTD) into a quasi-rigid unit, which separates the S6 four-helix bundle and opens the channel. Further out-of-plane rotation of the quasi-rigid unit pushes S6 towards the central axis, closing (inactivating) the channel. The inactivated conformation is characterized by a downward conformation of the cytoplasmic assembly, a tightly-knit subunit interface contributed by a fully occupied and partially remodeled Ca^2+^ activation site, and two salt bridges between the EF hand domain and the S2-S3 loop of the neighboring subunit validated by naturally-occurring disease-causing mutations. Ca^2+^ also bound to ATP, mediating a tighter interaction between S6 and CTD. Our study suggests that the *closed-inactivated* is a distinctive state of the RyR1 and its transition to the *closed-activable* state is not a simple reverse of the Ca^2+^ mediated activation pathway.

## Introduction

Ryanodine Receptors (RyRs) are complex intracellular, multi-domain Ca^2+^ channels that generate Ca^2+^transients in cells of higher metazoans. They are pivotal for contraction of skeletal and cardiac muscles (Flucher & Franzini-Armstrong, 1996, Meissner, 2017, Ríos, 2018), supporting transient rise in cytosolic Ca^2+^ from its resting level of ~100 nM, which is driven by a more than 1,000 fold Ca^2+^ concentration gradient across the membrane of the endoplasmic reticulum (sarcoplasmic reticulum in muscle, SR) (Ziman *et al*, 2010). RyRs also play a role in neuron excitability (Albrecht *et al*, 2001, Arias-Cavieres *et al*, 2018, Bouchard *et al*, 2003) and a myriad of other Ca^2+^ dependent pathways such as differentiation, survival and apoptosis (Bagur & Hajnóczky, 2017, Tu *et al*, 2016). Dysregulation of the channel leads to several life-threating diseases such as malignant hyperthermia, central core disease, sudden cardiac death, and lethal fetal akinesia deformation sequence syndrome (Alkhunaizi *et al*, 2019, McCarthy *et al*, 2000, Priori *et al*, 2001, Tiso *et al*, 2001, Treves *et al*, 2008).

The high-conductance RyR channel is strongly regulated, with cytosolic Ca^2+^ concentration having a biphasic effect on the probability of channel opening. Notably, both intracellular Ca^2+^ channels, RyRs and inositol P_3_ (IP3) receptors, display a bell-shaped curve of Ca^2+^ dependence (Bezprozvanny *et al*, 1991, Meissner, 2017, Meissner *et al*, 1986), characteristic of dual regulation by cytosolic Ca^2+^. In RyR, the ascendant branch of Ca^2+^ dependence with half-maximal concentration (*K_a_*) of 1-5 μM (Laver, 2018, Laver & Lamb, 1998) is mediated by a high affinity cytosolic Ca^2+^ site (des Georges *et al*, 2016). Ca^2+^ concentrations above 100 μM, reached in the tight nanodomain surrounding the RyR upon its opening (Langer & Peskoff, 1996), inhibit the channel (Chen *et al*, 1997, Xu & Meissner, 1998, Yamaguchi, 2020) which rapidly enters a refractory period that has been proposed to prevent depletion of the SR (Bers, 2002, Ríos *et al*, 2008). Out of the three isoforms, the “skeletal muscle” RyR1 isoform studied here is the most sensitive to Ca^2+^ inactivation (Meissner, 2017), where inactivation-impairing mutations are known to cause malignant hyperthermia (Gomez *et al*, 2016). While inactivation of RyR by Ca^2+^ has been characterized at the functional level, its underlying mechanism is unknown. Outstanding questions are whether high Ca^2+^ induces an allosteric change in RyR that dampens the affinity of its activation site for Ca^2+^, and whether the Ca^2+^ inactivated conformation is similar or distinct with respect to the closed conformation obtained in the absence of Ca^2+^.

Here, cryo-EM and single-particle 3D reconstruction were applied to rabbit RyR1 embedded in nanodiscs under conditions of partial Ca^2+^ inactivation. Classification of two independent cryo-EM datasets revealed, in both cases, coexistence of closed and open conformations, in agreement with functional experiments performed on the same channels using tritiated ryanodine binding. In the closed (inactivated) conformation, the resolution of the cryo-EM maps of nanodisc-embedded RyR1 enabled visualization of two lipids buried in a pocket of the transmembrane domain (TMD). To our knowledge, this is the first time that lipid is visualized in direct contact with the RyR. The open state, obtained by classification of the same dataset, represents the first RyR1 open conformation achieved in the absence of any extra activator other than the physiological activators, Ca^2+^ and ATP. We also carried out a control 3D reconstruction of RyR1 under the same conditions except for the absence of Ca^2+^, which yielded a closed channel. Comparison of the closed conformations of RyR1 at high Ca^2+^ and RyR1 in the absence of Ca^2+^ revealed unique features associated to Ca^2+^-inactivation. Both Ca^2+^-induced activation and inactivation can be explained by a unifying mechanism that involves conformational rearrangements within the central region of RyR1. Thus, the 3D structures of closed, open and inactivated RyR1 embedded in lipidic nanodisc provide a mechanistic framework to understand the biphasic response of RyR1 to Ca^2+^. In addition, two inter-subunit salt bridges appear to mediate the Ca^2+^-inactivated structural rearrangement, a finding supported by disease-causing mutations hindering such interactions and known to impair Ca^2+^-induced inactivation of the RyR1.

## Results

### Experimental design and functional validation of RyR1

Experimental conditions were fine-tuned to obtain the inactivated state. The channels were prepared in 2 mM free Ca^2+^, a concentration that inactivates the channel. As RyR1 is constitutively bound to the sensitizing ATP in the muscle cell (Kushmerick *et al*, 1992, Meissner et al., 1986, Xu *et al*, 1996), we included its non-hydrolyzable form, AMP-PCP (ACP) for its structural determination. In order to visualize the direct effect of Ca^2+^ on RyR1’s conformation FKBP12, a stabilizer of the cytoplasmic domain, was excluded. To determine the structure of RyR1 in a detergent-free,- membrane-embedded natural state, RyR1 was reconstituted into membrane scaffold protein (MSP) 1E3D1 nanodiscs in the presence of phosphatidylcholine, an abundant phospholipid in membrane fraction preparations. As a control, we carried out cryo-EM and 3D reconstruction of RyR1 using identical buffer conditions and saturating ACP, and substituted Ca^2+^ by 1 mM EGTA plus 1 mM EDTA (dataset denominated RyR1-ACP/EGTA). The channel was also reconstituted into nanodiscs in the presence of phosphatidylcholine.

The Ca^2+^-induced activity profile of rabbit RyR1 in SR membranes was determined using the tritiated ryanodine binding assay, which reflects the probability of channel opening. The assay indicated RyR1 channel inactivation at above 0.1 mM Ca^2+^, with an IC_50_ of 0.6 mM (Fig 1a). Presence of 2 mM ATP increased the efficacy of Ca^2+^-induced activation by ~3-fold at peak activation with similar potency for Ca^2+^-induced inactivation (IC50 of 0.7 mM), in agreement with earlier results obtained in lipid bilayer (Laver *et al*, 1995). We replicated the ryanodine binding experiments over the same Ca^2+^ concentration range with 2 mM ACP. Maximal ryanodine binding was ~1.5 fold higher in the presence of ACP compared to ATP, and in this case slightly higher Ca^2+^ concentration was necessary for the same degree of RyR1 inhibition, with an IC50 value for Ca^2+^ of 1.5 mM in the presence of ACP. Thus, under our experimental conditions of 2 mM free Ca^2+^, RyR1 inactivation was incomplete.

**Figure 1.**
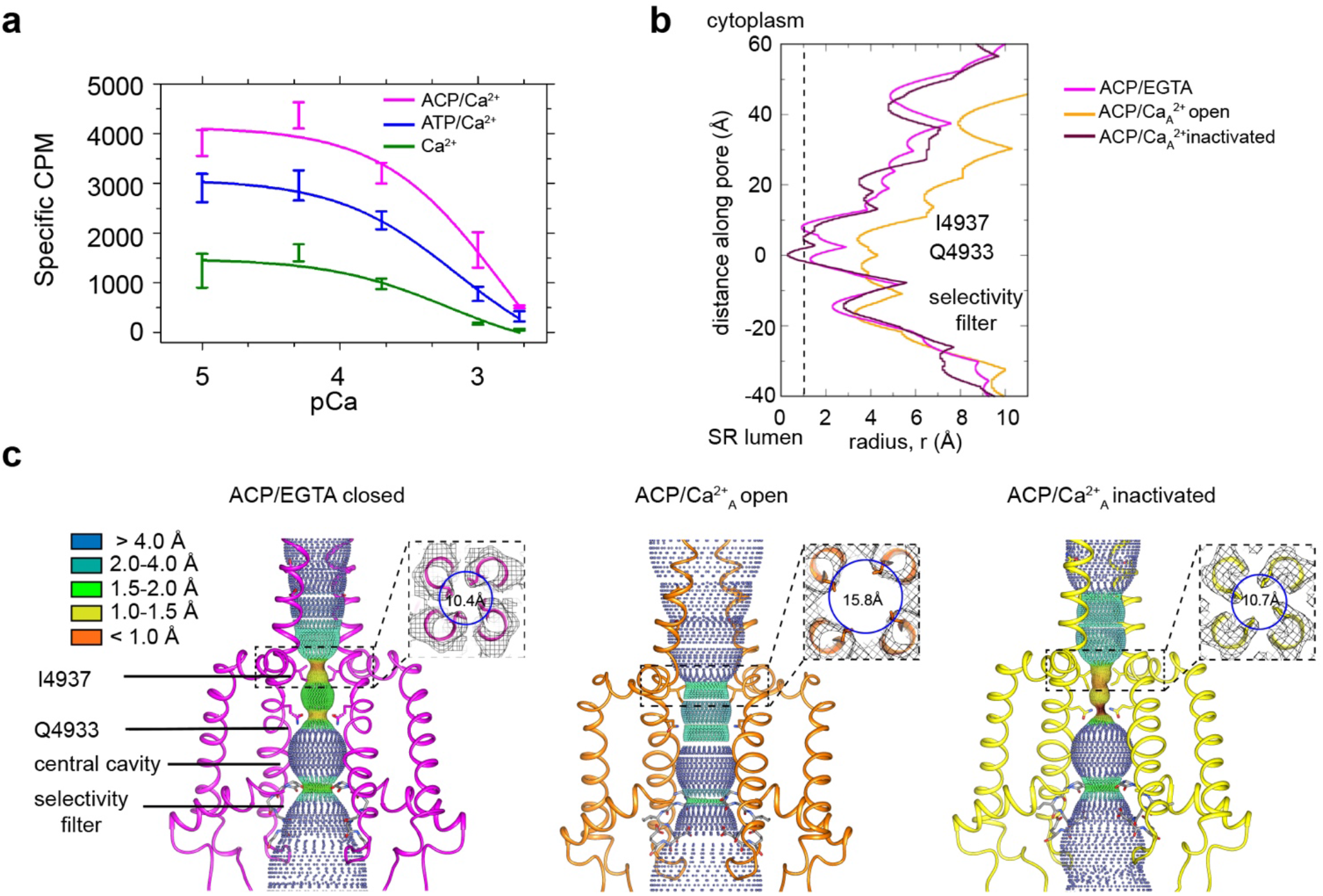
RyR1 at high Ca^2+^ concentration exhibits a mixture of open and inactivated conformations. **a,** Ryanodine binding of rabbit skeletal SR microsomes showing the Ca^2+^-induced inactivation of RyR1; activators ATP or ACP (2 mM) increased channel open probability by 3- and 4.5-fold, respectively. Ca^2+^ concentrations of 100 μM −2 mM progressively decrease the probability of opening (Po). Mean specific [^3^H]-ryanodine binding ± SE from four independent experiments. **b,**Pore profile of RyR1-ACP/EGTA, and of open and inactivated conformations in the RyR1-ACP/Ca^2+^_A_ dataset calculated with the program HOLE (Smart et al., 1993). The position of relevant landmarks is indicated. Radius corresponding to a dehydrated Ca^2+^ ion shown with a dashed line. **c,**Dotted surfaces of RyR1 ion permeation pathway in closed, open and inactivated conformations, color-coded according to pore radius. Overlaid coordinates of the S5, S6, pore helices and selectivity filter are shown for two diagonal protomers. Insets, cytoplasmic views of the Ile4937 constriction and corresponding pore diameter measured at the Cα backbone. Similar results were obtained for RyR1-ACP/Ca^2+^_B_ (Fig S5).

### Classification of the high Ca^2+^ dataset reveals an *open* conformation and a *closed-inactivated* conformation

Cryo-EM of rabbit RyR1 embedded in nanodisc in the presence of 2 mM ACP and 2 mM free Ca^2+^ followed by single-particle analysis and 3D classification revealed two classes of particles, with their pore either fully open or fully closed. The findings, reproduced in two independent datasets (A and B), reflected the partial inhibition observed at 2 mM free Ca^2+^ (pCa 2.7) in the ryanodine binding studies (Fig 1a). The 3D reconstructions of RyR1-ACP/Ca^2+^ open reported here are the first RyR1 open structures obtained in the presence of natural activators alone, probably owing to the higher free Ca^2+^concentration used as compared to previous studies. The RyR1-ACP/Ca^2+^ reconstructions with a closed pore suggest a distinct conformation that we refer to as RyR1-ACP/Ca^2+^-inactivated henceforth, as further supported by our data.

A symmetry of C4 was imposed after ascertaining true four-fold symmetry of the protein. The closed-pore (inactivated) channels were resolved to 3.8 Å and 4.3 Å resolution for the A and B datasets, respectively, which improved to 3.2 Å and 3.4 Å respectively using a phase improvement procedure (Terwilliger *et al*, 2020). Despite this improvement, we followed a conservative approach and used the non-density modified maps for subsequent analysis. The open conformation represented a smaller fraction of the data (13% and 19% for the A and B datasets), which limited the resolution to 4.6 Å and 5.8 Å, respectively (Figs S1-S3). The RyR1-ACP/Ca^2+^A open subset improved to a resolution of 4.0 Å after a symmetry expansion step. Unless specified, the reported observations correspond to dataset A.

Classification of the control RyR1-ACP/EGTA dataset yielded a single class with resolution of 4.3 Å (0.143 FSC gold standard), but further symmetry expansion and focused classification using a quadrant-shaped mask increased resolution to 3.9 Å (Fig S4). Despite the presence of the ACP activator, the 3D reconstruction of RyR1-ACP/EGTA yielded a closed state. Tables S1 and S2 summarize the cryo-EM data collection, single-particle image processing, and model quality attributes for all datasets, and Table 1 complies the main characteristics of the three conformations analyzed (closed, open, inactivated).

**Table 1.**
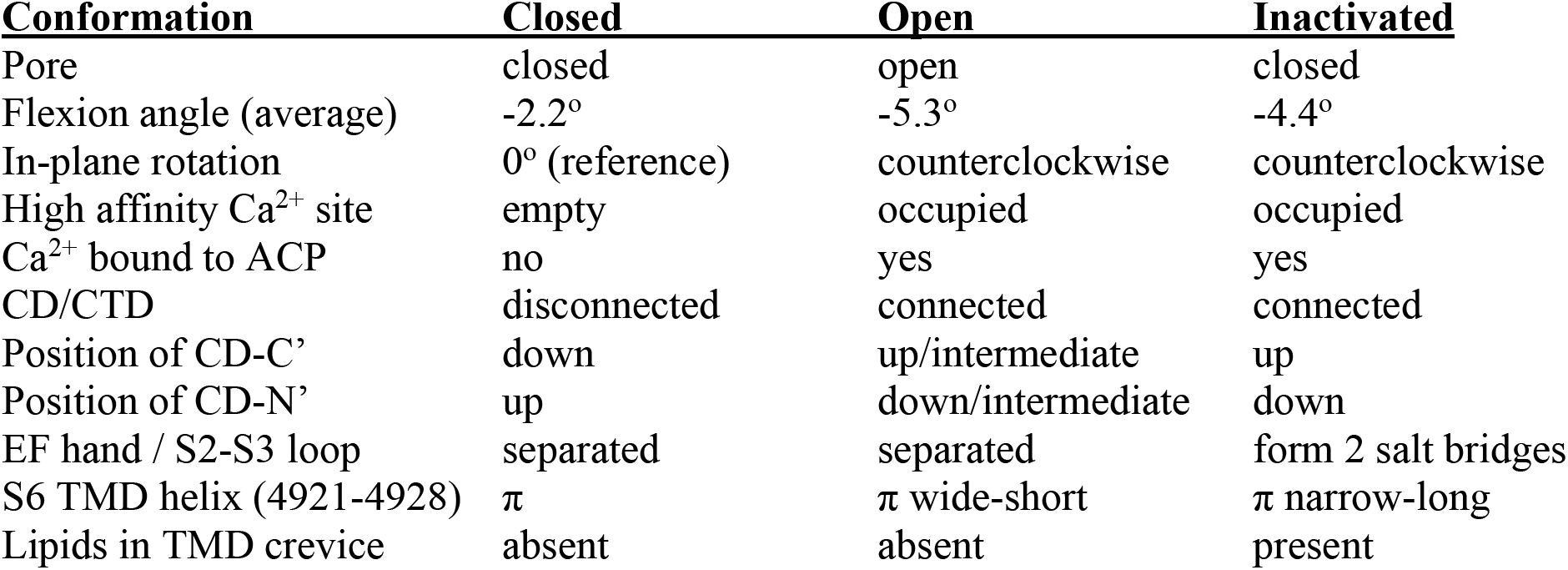
Differences between closed, open and inactivated conformations.

Pore analysis using the program HOLE (Fig 1b, c) of RyR1-ACP/EGTA indicated a closed pore with a radius constricting to 1 Å at the known hydrophobic gate Ile4937. In both RyR1-ACP/Ca^2+^ open A and B datasets the hydrophobic gate as appraised by the pore profile (Fig 1b, Fig S5) and as measured at the S6 helix backbone C_α_ atoms (for more accurate measurement according to the lower resolution of the two datasets) (Fig 1c, Fig S5) had diameters of 15.8 Å and 16.0 Å, which indicates an open, Ca^2+^-permeable pore in both cases. Pore dimensions are comparable to the pore diameter of rabbit RyR1 open structures obtained in the presence of activators such as Ca^2+^/PCB95 or Ca^2+^/Caffeine/ATP (16.7 Å; PDB ID: 5TAL (des Georges et al., 2016)). The slightly narrower pore in our case could be attributed to the millimolar instead of submicromolar Ca^2+^ in our case, and/or extra activators in addition to endogenous physiological activators in earlier open structures.

The reconstructions corresponding to RyR1-ACP/Ca^2+^ inactivated of the A and B datasets displayed a pore diameter at the Ile4937 gate of ~2 Å when considering the side chains, and 10.7 Å and 11.3 Å diameter respectively when measured at the backbone C_α_ atoms, making it impermeable to Ca^2+^ ions (Fig 1b, c, Fig S5). The position of the S6 backbone at Ile4937 is similar to our closed structure (r=10.4 Å) and to RyR1-FKBP12/EGTA (r=10.3 Å; PDB ID: 5TB0 (des Georges et al., 2016). Interestingly, the channel pore was narrower than 1 Å at Gln4933 in RyR1-ACP/Ca^2+^_A_ inactivated (Fig 1b, c).

We previously established that the large square-shaped cytoplasmic shell of the RyR undergoes a conformational change upon opening. The periphery of its four quadrants tilts downwards towards the membrane, while their inner corner tilts away from it, rotating around a pivot point (Samsó *et al*, 2009). Such tilt can be quantified with the flexion angle, whereby negative angle (downward) correlates with opening, positive angle with closing, and absence of FKBP lowers the flexion angle of closed states (Steele & Samsó, 2019). In general, the approximate ranges of flexion angles are +1° to +2° for RyR1-FKBP12/EGTA closed, −1° to −3° for RyR1-EGTA closed, 0° to −2° for RyR1-FKBP12 primed (with activating ligands and closed pore), and −1.5° to −5° for RyR1-Ca^2+^ open with or without FKBP12 (Iyer *et al*, 2020, Steele & Samsó, 2019). Here, RyR1-ACP/Ca^2+^ open channels have flexion angles of −5.1° and −4.8° for the A and B datasets, within the expected range. The consensus 3D reconstruction of RyR1-ACP/EGTA had flexion angle of −2.2°, and its two classes had flexion angles of - 1.8° (24%) and −2.9° (76%) (Fig S4). But unexpectedly for closed-pore channels, RyR1-ACP/Ca^2+^ inactivated datasets have a negative flexion angle (−4.6° and −4.2° for A and B datasets, respectively) (Figs S1, S3). As there was indication of variability in the cytoplasmic shell of RyR1-ACP/Ca^2+^A inactivated dataset, we carried out fourfold symmetry expansion and focused classification for the monomer and obtained three sub-classes, all showing downward motion: class 1 (22 % of particles, 3.1 Å resolution, −3.6° flexion angle), class 2 (44 % of particles, 2.9 Å resolution, −4.4° flexion angle) and class 3 (20 % of particles, 3.3 Å resolution, −5.6° flexion angle) (Fig S1). Such pronounced downward flexion angles in a closed channel cannot be explained just by the lack of FKBP12 in our preparations. The distinct conformation of RyR1-ACP/Ca^2+^inactivated, consisting of a closed pore and extreme-downward cytoplasmic assembly, prompted further analysis of the central region that joins the cytoplasmic and transmembrane domains.

### Different arrangement of the central region in Ca^2+^-inactivated, closed and open conformations

In RyR1-ACP/Ca^2+^ open, the high-affinity Ca^2+^ binding site formed by Glu3967, Glu3893 (from the CD; residues 3668-4070), and Thr5001 (from the CTD; residues 4957-5037) as reported earlier (des Georges et al., 2016), and additionally Gln3970 in our case, was visible up to 20 σ. The Ca^2+^-induced reorientation of CTD with respect to CD and subsequent separation of S6 was obvious when compared to the RyR1-ACP/EGTA structure (Fig 2a, Fig S6), consistent with previous reports of Ca^2+^-induced activation (Bai *et al*, 2016, des Georges et al., 2016).

**Figure 2.**
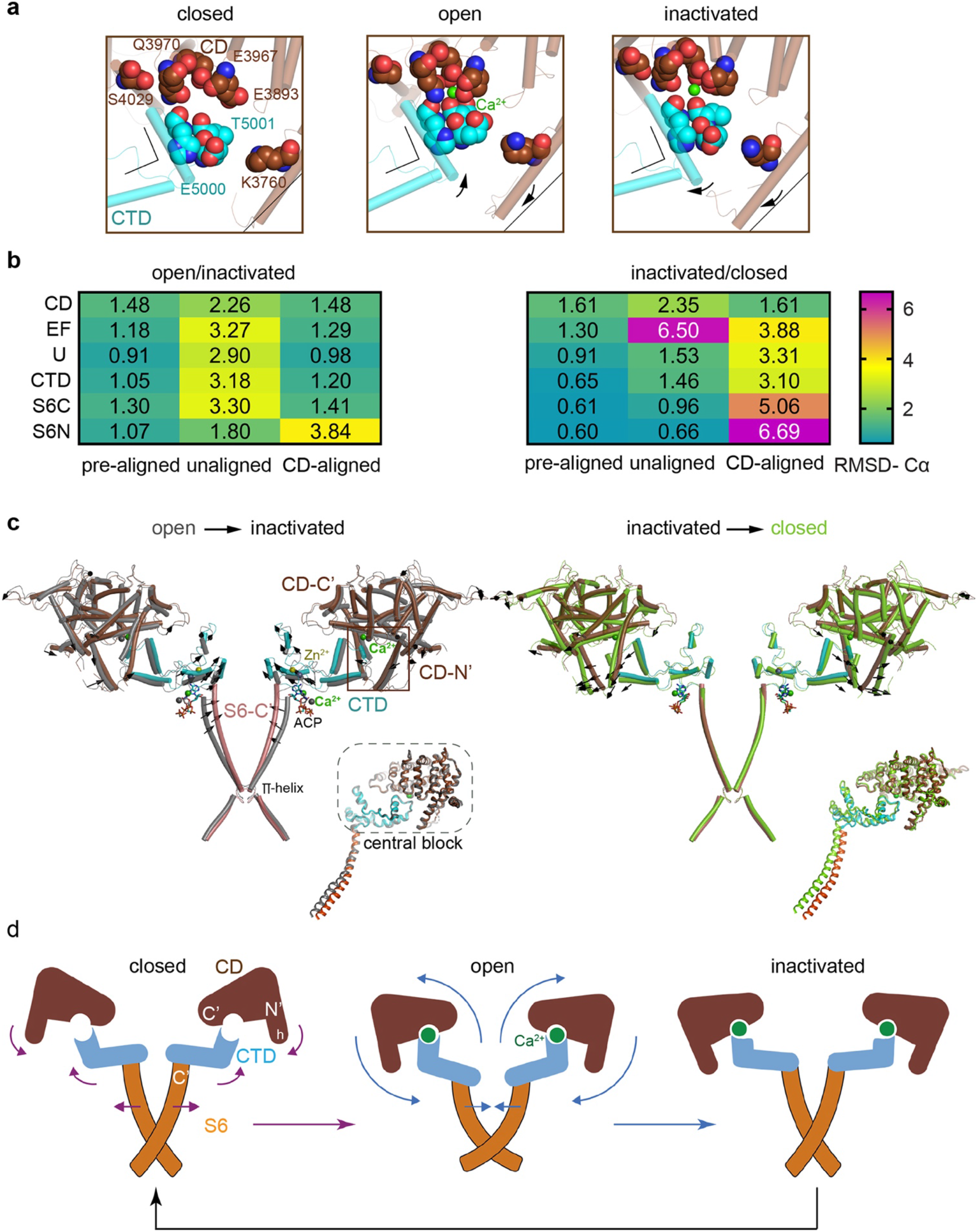
Inactivation of RyR1 involves out-of-plane rotation of the central block and rearrangement around the Ca^2+^ activation site. **a,** The Ca^2+^ binding site in the CD/CTD interface in different conformations with involved residues shown with spheres. Residues Glu3983, Glu3967, Gln3970 (CD), and Thr5001 (CTD), within 2.8 Å from Ca^2+^ in the open state, are shown. In RyR1-ACP/Ca^2+^ inactivated, Gln3970 switches its interaction to Ser4029 (contact within 3.6 Å). Arrows and stationary reference lines illustrate the conformational changes undergone with respect to the panel on the left. The region represented relative to the channel is highlighted with a square in panel c. See Fig S6 for the corresponding cryo-EM densities. **b,** Heat map showing Cα-backbone RMSD (in Å) between domain pairs from respective conformations after aligning them (pre-aligned), in their native conformation (unaligned), and after aligning the protomers through their respective CD (CD-aligned). Their difference represents the RMSD change caused by domain relocation. **c,** Overlaid structures of the CD-CTD-S6 domains in different conformations; only two protomers shown for clarity. Left, in the transition from RyR1-ACP/Ca^2+^ open (gray) to RyR1-ACP/Ca^2+^ inactivated (colored), the CD-CTD block, “connected” by Ca^2+^ coordination, undergoes an out-of-plane rotation around the Ca^2+^ binding site that pushes S6C’ towards the pore axis, closing the channel. Right, in the transition from RyR1-ACP/Ca^2+^ inactivated (colored) to RyR1-ACP/EGTA (green), Ca^2+^ unbinding disconnects the CD from the CTD. Structures at the bottom right of each panel show the comparison of the CD-CTD-S6C’ of the central block after forcing superimposition of their respective CDs. **d,** Schematics of the conformational changes from RyR1-ACP/EGTA (closed), to RyR1-ACP/Ca^2+^ open, to RyR1-ACP/Ca^2+^ inactivated. Activation, where Ca^2+^ binds to the high affinity site, is required prior to inactivation. The CD-protruding fourth helix (3753-3769) is indicated by “h”. Colored arrows indicate the conformational change towards the structure on the right.

An outstanding question is how higher Ca^2+^ concentration may result in inactivation. Potential mechanisms are an additional allosteric change that dampens the affinity of the Ca^2+^ activation site, or that the high affinity Ca^2+^ site remains occupied while an additional conformational change overcomes activation. Consistent with the second scenario, the high affinity Ca^2+^ binding site in RyR1-ACP/Ca^2+^ inactivated remained fully occupied by Ca^2+^ up to 20 σ, although with additional out-of-plane tilting of the CD and CTD around the Ca^2+^ binding site. The Ca^2+^ ion was coordinated by Glu3967, Glu3893 (CD), and Thr5001 (CTD) within contact distances of 2.8 Å, consistent with the open structures. However, on the CD-C’ side, Gln3970 lost coordination to Ca^2+^ (when compared to RyR1-ACP/Ca^2+^ open) interacting with Ser4029 instead, while on the opposite CD-N’ side an additional contact formed between residues Lys3760 located in the protruding fourth helix of the CD (helix “h”; residues 3753-3769) and Glu5000 (Fig 2a, Fig S6). Mutations Q3970E/K in RyR1 are implicated in central core disease and equivalent RyR2 mutations in cardiac arrythmia (Chirasani *et al*, 2019), which highlights the important Ca^2+^ sensing role for this residue.

To understand how the fully occupied high affinity Ca^2+^ binding site (CD/CTD interphase) led to a closed pore in RyR1-ACP/Ca^2+^ inactivated, we looked for differences between open and inactivated in the domains spanning from the CD/CTD interphase to the pore. Specifically, we focused on domains around the high affinity Ca^2+^ binding site (CD, residues 3668-4070; EF hand, residues 4071-4131; U-motif, residues 4132-4251; and CTD, residues 4957-5037) as well as the N’ and C’ sections of S6 (S6N’ and S6C’). Comparison was done for dataset A after confirming similarity for the encompassed 672 residues of 0.95 Å Cα root mean square deviation (RMSD) between inactivated A and B datasets (Fig S7a). The RMSD (Cα backbone) between domains from RyR1-ACP/Ca^2+^ inactivated versus open was below 1.5 Å for all comparisons when the domains were pre-aligned (Fig 2b, Fig S7b, c), indicating little conformational change. Without pre-alignment, RMSD was higher (Fig 2b), indicating domain repositioning, the degree of which was estimated by subtracting the “pre-aligned” from the “unaligned” RMSDs. This yielded an average shift of ~2 Å per domain in going from open to inactivated, except for S6N’ where the shift was ~0.7 Å (Fig 2b). Comparing the same domains between RyR1-ACP/Ca^2+^inactivated and RyR1-ACP/EGTA (closed) yielded smaller repositioning with average shifts of ~1 Å, reflecting similarity in the domain’s locations, with the exception of the EF hand domain that undergoes a ~5 Å repositioning (Fig 2b, Fig S7c).

When the entire 672 residue-block were pre-aligned using the CD domain (CD-aligned), there was good overlap between open and inactivated (RMSD below 1.5 Å) except for S6N’ (3.8 Å), revealing the bend in the middle of S6 upon opening. In contrast, CD-aligned RMSD of inactivated vs closed is higher, between 3-4 Å for the EF hands, U-motif and CTD, and increases towards the C-terminus, 5 Å for S6C’ and almost 7 Å for S6N’ (Fig 2b, “CD-aligned”). This indicated that domains CD, EF hands, U-motif, S6C’, and CTD relocated together forming a rigid central block in the presence of Ca^2+^ acting at the CD/CTD interface. Taken together, these results imply that conformational changes from CD are transmitted to S6, only in the presence of Ca^2+^ (Fig 2c, d).

We analyzed how the central block of 672 residues may evolve from RyR1-ACP/EGTA (closed), to RyR1-ACP/Ca^2+^ open, to RyR1-ACP/Ca^2+^ inactivated. Upon Ca^2+^-induced opening, “engagement” of CD and CTD by Ca^2+^ tilts the CD out-of-plane (such that helix h tilts 3° inwards), while the CTD tilts up and outwards, separating the S6 helices directly connected to that domain (Fig 2c, d). To reach the inactivated state, the central block tilts further (see rotation axis in Fig S7b) in a movement similar to pushing down the levers of a winged corkscrew, which pushes each S6 helix 2.5 Å towards the pore axis, closing the channel. Alpha helices within the CTD act as a lever coupling this out-of-plane rotation to S6C’; the conformational change is more clearly seen in the schematics in Fig 2d and Movie S1. As described further below, ATP reinforces the connection between CTD-S6C’ in the presence of Ca^2+^.

### Formation of salt bridges between EF hand domain and S2-S3 loop of the neighboring subunit in the inactivated state

The EF hand domain of RyR1, a candidate Ca^2+^ regulation site (Du *et al*, 1998, Gomez et al., 2016, Gomez & Yamaguchi, 2014, Xiong *et al*, 1998), did not show significant changes, with maximum Cα RMSD of 1.5 Å among the different states analyzed (closed, open, inactivated). The EF hand loops (residues 4081-4090, 4116-4123) were empty in our high Ca^2+^ conditions, which is consistent with the low affinity of Ca^2+^ to this site (Kd 3.7 mM (Xiong et al., 1998)). Nonetheless, the entire EF hand domain, which protrudes from the CD-C’, repositions noticeably during activation, with a 3.4° counterclockwise in-plane rotation (as seen from the cytoplasmic side), and 6.8° upward out-of-plane rotation. This movement brings the EF hand domain in closer proximity to the S2-S3 loop (residues 4664-4786; cytoplasmic loop between S2 and S3 TMD helices) (Fig 3). With inactivation, an additional 1° counterclockwise rotation and 2.7° upward out-of-plane rotations define a physical limit to the counterclockwise motion, forming two salt bridges at this inter-subunit interface: Glu4075-Arg4736 and Lys4101-Asp4730, with their side chains within 3.5 Å (Fig 3a). This interaction is critical to support inactivation, as demonstrated by the fact that MH/CCD mutations facing this interface F4732D, G4733E and R4736W/Q, the latter including the Arg directly involved in the salt bridge (see Fig S3b), greatly reduced channel inactivation (Gomez et al., 2016). On the other hand, MH/CCD mutations T4082M, S4113L, and N4120Y, in regions of the EF hand domain away from the interface (Fig S3b), did not affect RyR1 inactivation (Gomez et al., 2016), serving as a negative control for this hypothesis.

**Figure 3.**
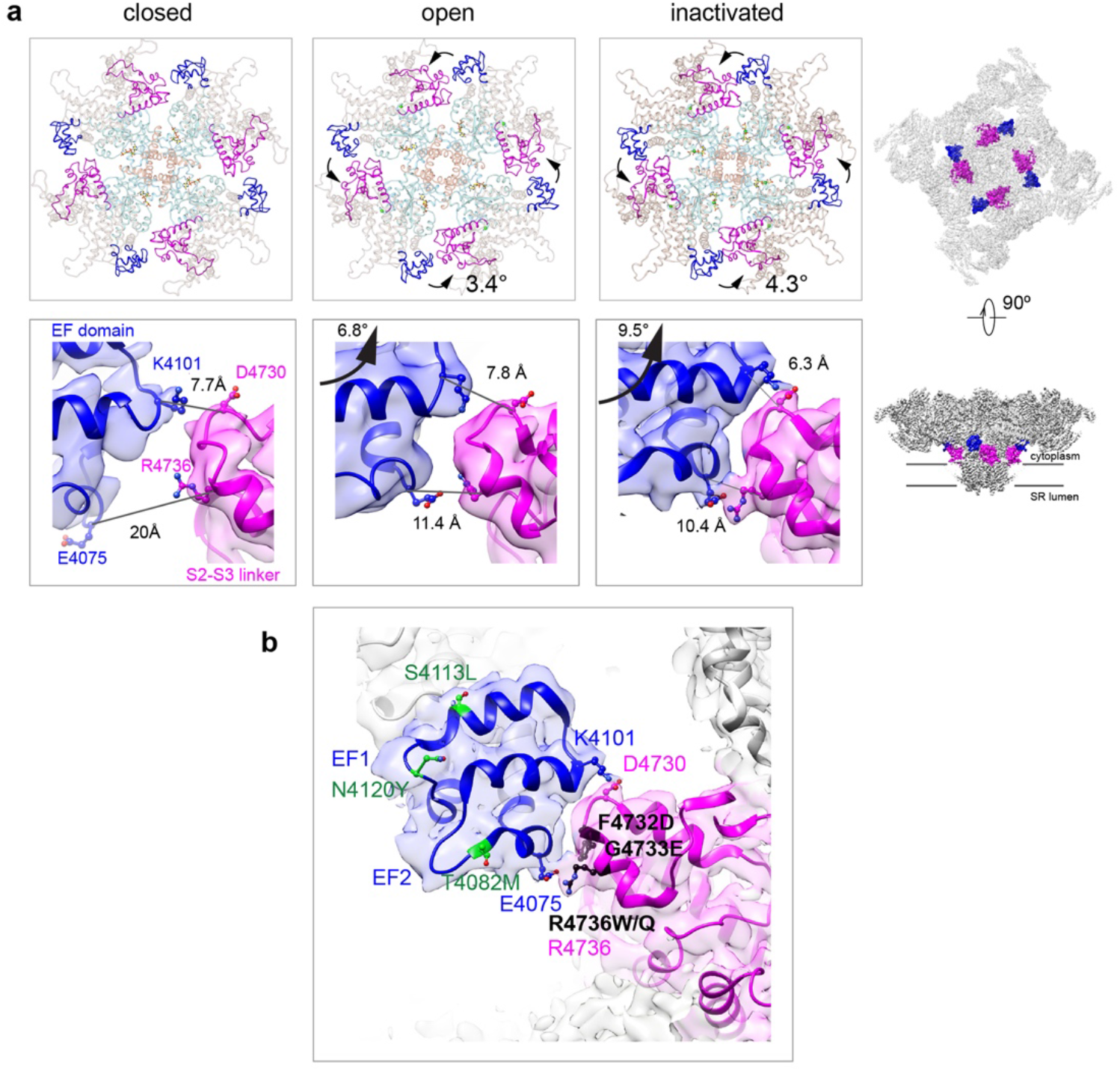
Two salt bridges between the EF hands and the S2-S3 loop are determinant for the inactivated state. **a,** Cytoplasmic SR view (top row) and side view (bottom row) highlighting the EF hand domain (blue) and S2-S3 loop (magenta) at the subunit interface. Counterclockwise in-plane and out-of-plane rotations of the central region in the transition from RyR1-ACP/EGTA (closed) to RyR1-ACP/Ca^2+^ open brings the two domains closer together. Further rotation of the CD (tan color in top row) and its protruding EF hand domain in RyR1-ACP/Ca^2+^ inactivated brings the two domains in contact. The inter Cα-Cα contact distances illustrate the progressive approximation of the two domains. Two salt bridges, Lys4101-Asp4730 and Glu4075-Arg4736 form in the RyR1-ACP/Ca^2+^ inactivated conformation. The location of the domains in the context of RyR1 is shown on the right panels. **b,** MH/CCD mutation sites in the interface between EF hand domain and S2-S3 loop that abolish Ca^2+^-dependent inactivation (black), versus MH/CCD mutations without effect on inactivation (green). The two EF loops are indicated. Residues forming the salt bridges are labeled according to domain color. R4736, directly forming the salt bridge, is susceptible to MH mutation.

### Ca^2+^ binds to the ATP binding pocket

The ATP binding site showed full occupancy in all conformations. Under closed-state conditions, ACP bound to the pocket formed by the U-motif, S6C’ and CTD (Fig 4a “closed”), in the same position as ATP (des Georges et al., 2016), with a high map significance (7σ). An additional elongated non-protein density was associated to ACP in the RyR1-ACP/Ca^2+^ open and inactivated maps. This distinct density has high map significance (15σ) (Fig 4a, Fig S8), suggestive of a putative Ca^2+^ ion. Well-resolved local density in RyR1-ACP/Ca^2+^A inactivated class-1 map allowed tentative modeling of two waters on either side of the putative Ca^2+^ that connect on either side to ACP’s γ-phosphate and Thr4979 (CTD) (Fig 4a “inactivated”), suggesting potential coordination through Ca^2+^’s first layer of hydration. Analysis of protein-ligand interactions based on the atomic coordinates (Laskowski & Swindells, 2011) showed an increased network of electrostatic and hydrophobic interactions, and three predicted hydrogen bonds, that the inactivated conformation gained with respect to closed conformation (Fig 4a). Thus, the nucleotide located deeper into the cavity in going from open to inactivated, increasing connectivity between S6C’ and CTD and reducing connectivity to the U-motif (Fig 4b). Interestingly, ACP in RyR1-ACP/Ca^2+^_A_ inactivated acquired an interaction with the backbone carbonyl of His4983, a residue that participates in the C2H2 Zinc motif which is central to the CTD (Fig 4). Based on the higher map significance of the putative Ca^2+^ ion in our cryo-EM maps obtained at higher Ca^2+^ concentration (Fig S8) and the enhanced inactivation when ATP is present (Sitsapesan & Williams, 2000), the interphase between the nucleotide, the CTD and S6 could be thought of as an atypical low affinity Ca^2+^ inactivation site. The picture is bound to be more complex, as presumably Mg^2+^ present in the cytoplasm would also bind to ATP. Thus, under physiological conditions of high local Ca^2+^, competition between the two divalent cations for ATP may take place.

**Figure 4.**
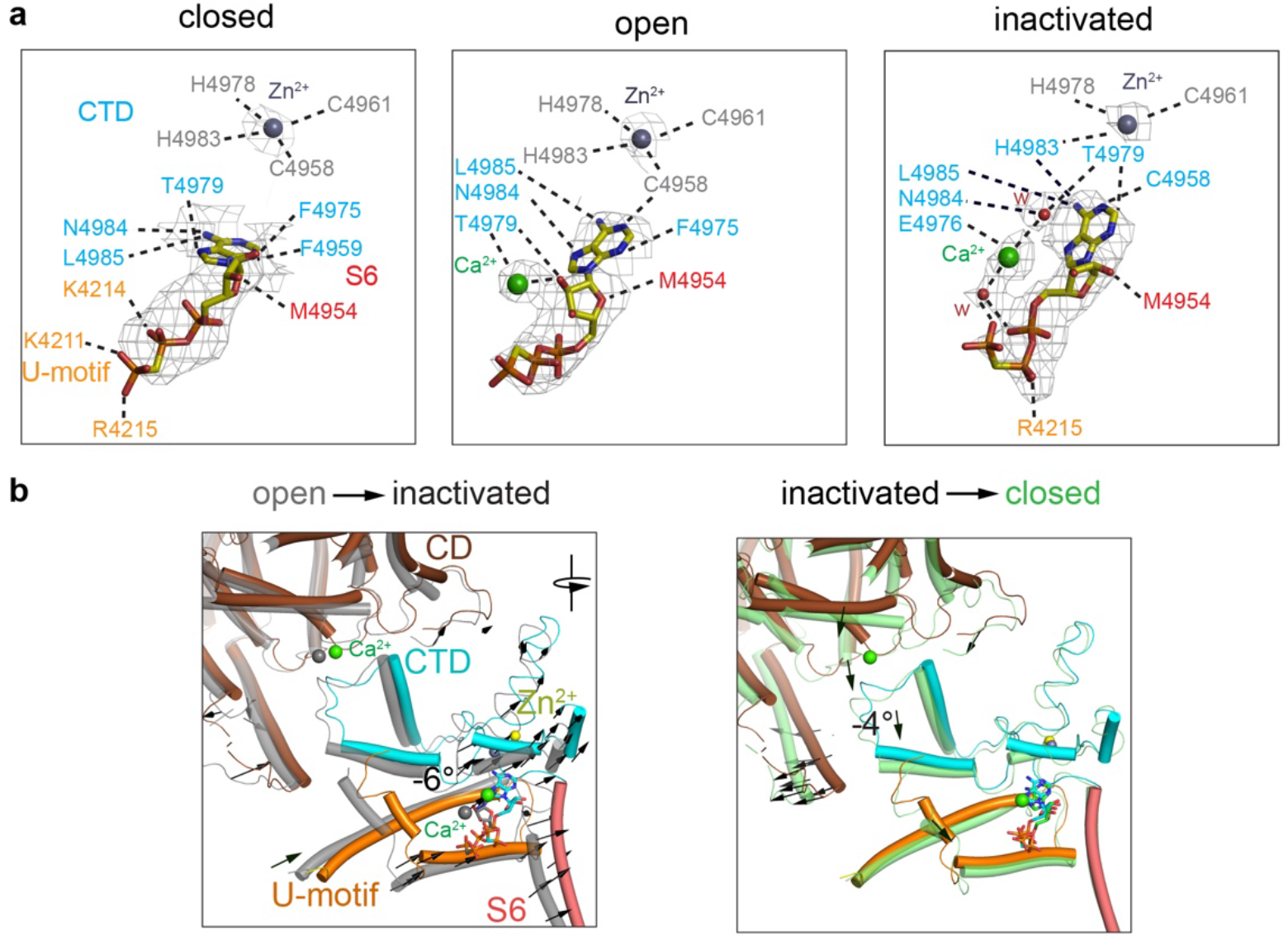
Changes in the interaction network of the nucleotide and complexed Ca^2+^. The ATP binding pocket is formed by the U-motif, CTD and S6C’. **a,** Cryo-EM densities (gray mesh) of ACP in the different states represented at map significance values (RMSD σ) above 12, 7 and 7, respectively. Residues within 4 Å of either ACP, Ca^2+^ or Zn^2+^ (from the CTD zinc finger) are color coded according to domain. Notably, an interaction between His4983 (CTD) and ACP is only observed in the inactivated state. Fitted water densities (w) are represented in red. **b,** Left, ACP goes deeper into the ATP binding pocket in RyR1-ACP/Ca^2+^ inactivated (colored structure, blue ACP) as compared to the open state (gray structure, red ACP) due to a 6° rotation (see arrows). Right, absence Ca^2+^ in the closed state (green) causes release of the CTD from the CD, resulting in a 4° tilt of the CTD and U-motif (see arrows). Additional reorganization allows closer interaction between ACP and U-motif in the closed state.

### Protein-lipid interactions within the nanodisc environment

To provide a more physiological environment to the TMD domain and avoid the presence of detergents, we embedded the protein in scaffold protein MSP1E3D1 that assembled into nanodiscs. The reconstituting lipid was phosphatidylcholine. Focused refinement of RyR1-ACP/Ca^2+^_A_ inactivated and open, resulting in reconstructions with 3.5 Å and 4.4 Å resolution respectively, resulted in a visible electron density for the nanodisc. Two molecules of the MSP1D1E3 scaffold protein wrapped closely around the TMD in a double-belt arrangement (Fig 5a). The larger top belt adopts a quatrefoil shape, with one voltage sensor (S1-S4) in each lobe, while the lower belt is rounder and smaller, following the tapered TMD. The larger footprint of the top half of RyR1’s TMD domain, reflected in the surrounding nanodisc, appears to correlate with the curvature of the membrane around RyR1 observed by electron tomography in its native membrane (Chen & Kudryashev, 2020, Renken *et al*, 2009). Even considering the tight fit between the nanodisc and RyR1’s TMD, conformational changes were unhindered and the channel opened within the nanodisc environment. The slight expansion of the top belt of the nanodisc at the level of the ion gate in the open channel (Movie S2) reveals a certain degree of plasticity of the scaffold protein.

**Figure 5.**
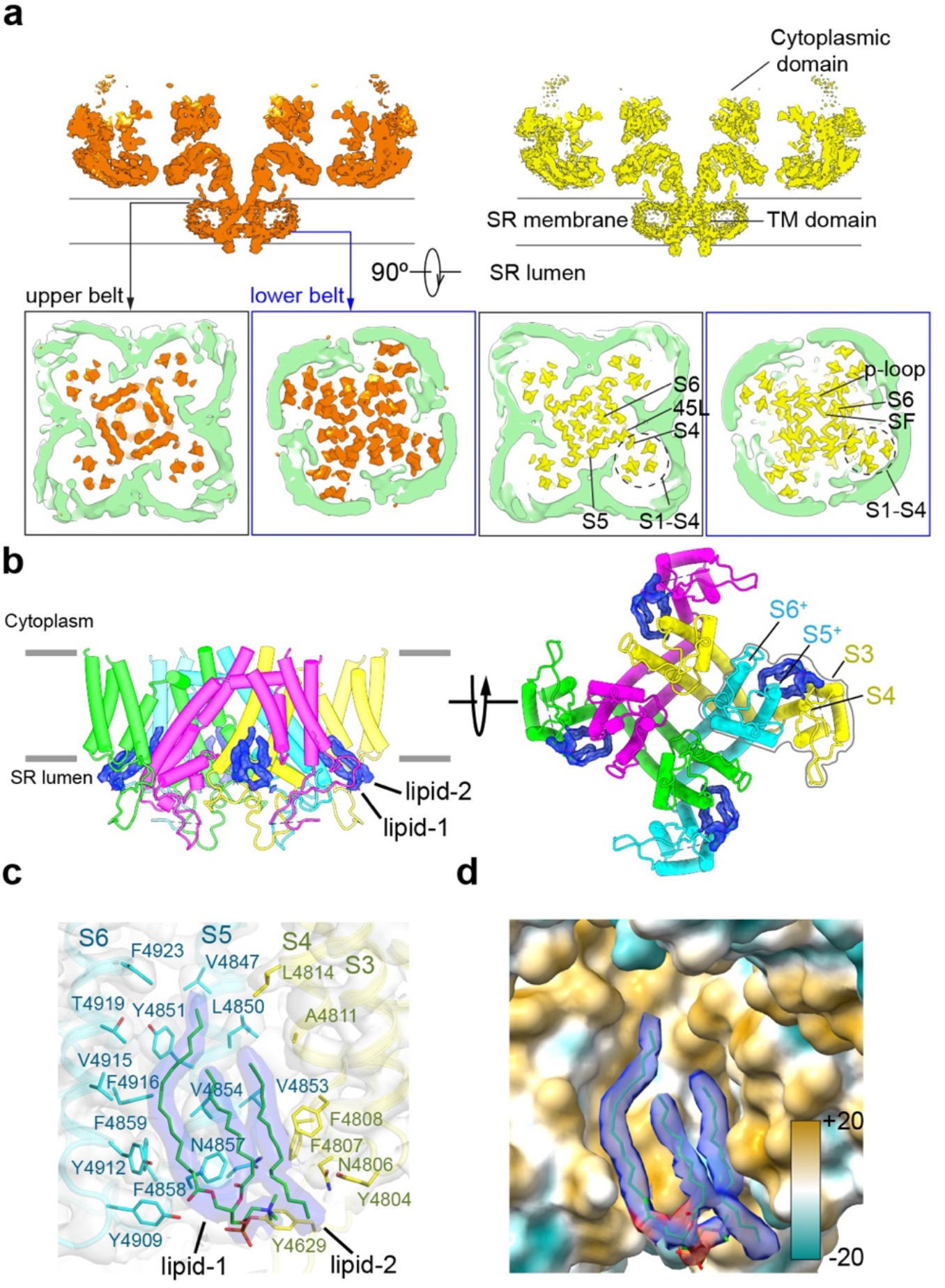
The nanodisc environment and visualization of lipids in a crevice of the TMD. **a,** Top, central slice of the side view of the RyR1-ACP/Ca^2+^_A_ open and RyR1-ACP/Ca^2+^_A_ inactivated cryo-EM maps highlighting the density corresponding to the upper and lower nanodisc belts. Bottom, corresponding views seen from the cytoplasmic direction. For clarity, the nanodisc density (green) was extracted and low-pass filtered to 7 Å resolution. The top belt of the nanodisc expands slightly to accommodate the conformational change; see Movie S2. **b,** Side and luminal views of the TMD of RyR1-ACP/Ca^2+^_A_ inactivated with putative lipid densities shown in blue. **c,** Lipid binding pocket of RyR1-ACP/Ca^2+^_A_ inactivated lined by lipophilic amino acids from S3 and S4 of the voltage sensor-like domain (S1-S4; yellow) and core helices S5 and S6 (cyan) from two different protomers. Amino acids within 5 Å from the lipids are shown (sticks) with their corresponding side chain densities. Electron densities corresponding to the lipids contoured at 8σ (in blue) are modeled as a PC (16:0-11:0) for lipid-1 and a 16 C acyl chain for lipid-2. **d,** Molecular lipophilicity potential of the surface lining the crevice, ranging from hydrophilic (cyan) to hydrophobic (golden). The hydrophobic tails of the lipids are shown in blue and the negative electrostatic surface potential of the polar lipid head is shown in red.

We resolved two lipid densities at the four inter-subunit interfaces in the RyR1-ACP/Ca^2+^_A_ inactivated CD-TMD focused map (Fig 5b). The two lipid densities, observed up to 8σ map significance, spanned ~20 Å across the inner leaflet of the SR membrane, at a hydrophobic pocket in the domain-swapped inter-subunit space formed by the S1-S4 bundle and core TMD helices (S5 and S6) (Fig 5b, c). One density (lipid-1), encompassing two fatty acyl tail moieties of 16 and 11 carbons with a polar head compatible with a phosphatidylcholine molecule was resolved near S5/S6 (Fig 5c). The lipid likely originates from a longer unsaturated PC (16:0-18:1) used in 50-fold molar excess while embedding RyR1 into nanodiscs. Lipid-2, with a resolved 16-carbon fatty acid tail, was sandwiched between lipid-1 and S3/S4. The lipid’s fatty-acyl tails interact extensively with 21 amino acids in the lipophilic pocket formed by S3/S4 and S5/S6 of neighboring subunits, while the density corresponding to the lipid head moiety was positioned within 5 Å from Tyr4629, Asn4857, and Tyr4909 (Fig 5c). Together, the lipids covered ~622 Å^2^ of the 1990 Å^2^ surface area of the S3/S4-S5/S6 subunit interface forming a hydrophobic crevice (Fig 5d). No lipid density was resolved in the RyR1-ACP/Ca^2+^_A_ open map: besides the lower resolution of this map, its crevice is narrower, as S3 and S4 remodeled the lipid binding pocket by tilting ~4.4° and ~3.9°, respectively. Furthermore, reorientation in Phe4808 (S4) and Tyr4912 (S6) in the open state would cause steric clash with lipid-2 and lipid-1 respectively (Fig S9), rearranging or even excluding the lipids in the open channel, a phenomenon similar to the observation reported for the TRPV3 channel (Singh *et al*, 2018).

### Plasticity of transmembrane helices and stability of the luminal mouth of the channel

The open and inactivated conformations obtained in the membrane-like environment provided by the nanodisc warranted a closer look at the TMD helices. We report perturbations in the alpha helical configuration, consisting of near-3_10_ helix in the N’ of S4 (residues 4807-4813) and π-helix in S5 (4856-4859) and S6 (4921-4928) (Fig 6a, b). These segments contain Phe clusters (FFF 4807-4809, FF 4858-4859, FFFF 4920-4923) conserved in all RyR isoforms and IP3R intracellular Ca^2+^ release channels; some of them lining the hydrophobic crevice and forming contacts with the lipids (see also Fig 5c).

**Figure 6.**
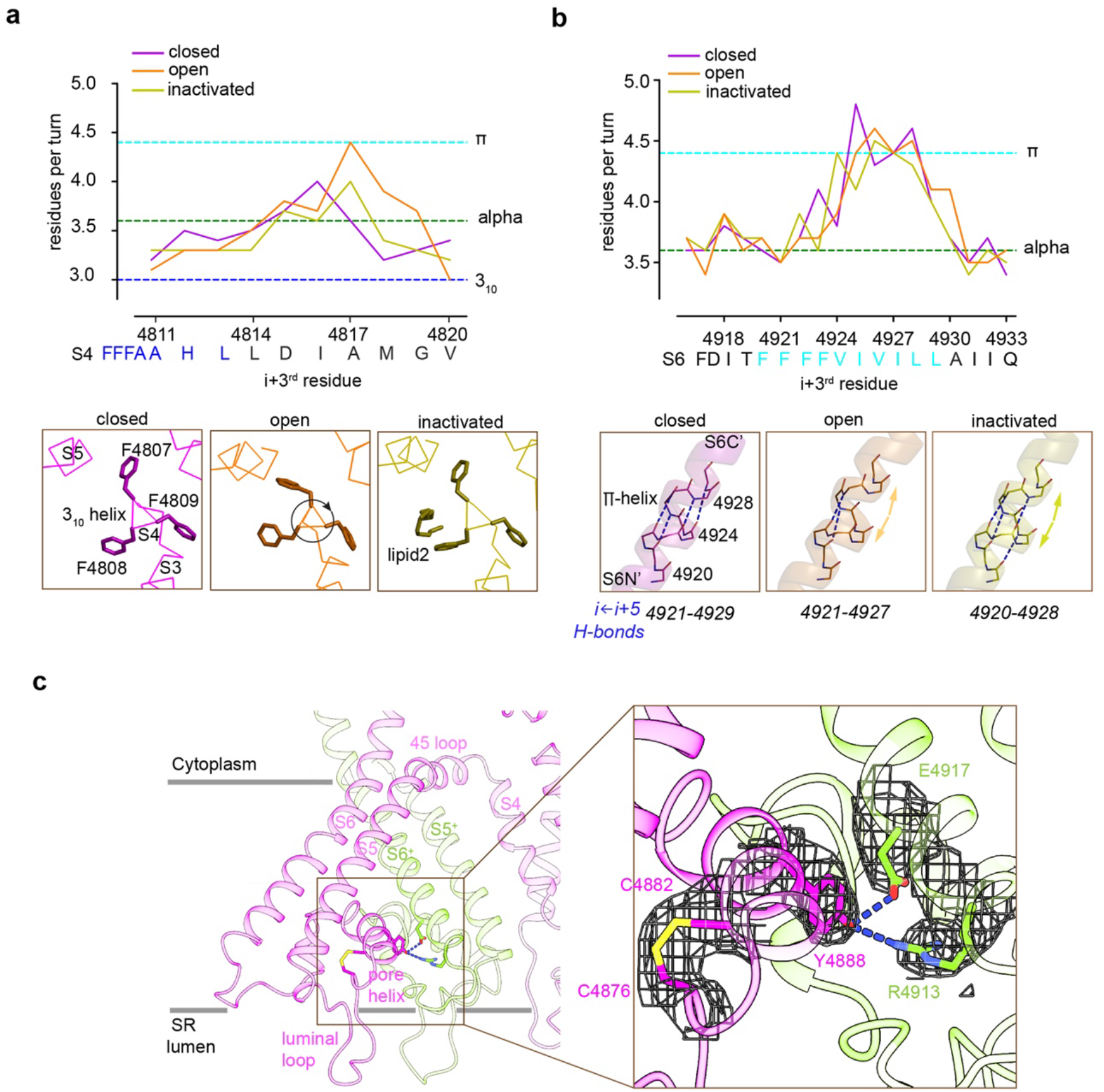
Local conformational changes in the transmembrane helices among closed, open and inactivated conformations. **a,** Top, residues per turn for S4 reveals a 3_10_ helix (4807-4813; blue dashed line) at its N’, and helical changes among the three conformations. Residues per turn are computed for a 3-residue moving window. Bottom, a segment of the S4 displaying the 3_10_ helix in FFF (4807-4809). Uncoiling of the S4-C’ in the open structure rotates the FFF region, including Phe4808 which interacts with lipid-2 in the inactivated structure. **b,** Top, the N’ region of S6 (residues 4920-4928) contains a tetra Phe motif (4920-4923) followed by a π-helix region. Bottom, the backbone hydrogen bonding network in the π-helix region in the three distinct conformations; dashed lines show unbroken i ← *i*+*5* hydrogen bonds. In this area, both open and inactivated channels lack hydrogen bonding which facilitates flexibility of the S6 helices. **c,** A Cys4876-Cys4882 disulfide bond formed between the luminal loop (4860-4878) and the pore helix (4879-4893) of the same subunit and hydrogen bonds between the pore helix and S6 from the neighboring subunit stabilize the luminal mouth of RyR1. The cryo-EM densities are shown in mesh.

In S4, comparison of the three conformations suggests that the degree of over-coiling of the N’ half of S4 is balanced by the uncoiling of its C’ half (residues 4814-4819) in the open structure (Fig 6a), which causes the reorientation of Phe4808 into the hydrophobic crevice. In S6, the π-helix region introduces non canonical i ←i+5 hydrogen bonding which is energetically less stable than a regular α-helix (Fodje & Al-Karadaghi, 2002, Kumar & Bansal, 2015)(Fig 6b). Both the inactivated and open structures lacked intra-helix hydrogen bonds within the π-helix (Fig 6b), which may facilitate the flexibility of S6 required for opening and inactivation. In RyR1, deletion of FF (4923-4924) in S6, which is associated to fetal akinesia deformation syndrome or FADS (Xu *et al*, 2020), has been shown to abolish Ca^2+^ activation, supporting a role for the Phe clusters and their structural transition in Ca^2+^-mediated gating.

The SR luminal loop (4860-4878) between S5 and the pore helix (4879-4893), proposed to act as luminal Ca^2+^ sensor (Sitsapesan & Williams, 1995) and as anchorage point for proteins within the SR lumen (Beard *et al*, 2009), contains six acidic residues (EDEDEPD 4867-4873). The loop is separated from the rest of the TMD and its acidic residues point to the lumen interior (Fig 6c). As our experimental high Ca^2+^ concentrations correspond to these in the SR lumen, the arrangement of the four luminal loops is probably a close representation of their native state. Our higher resolution RyR1-ACP/Ca^2+^ inactivated map allows to discern a disulfide bond (Cys4876-Cys4882) between the S5 and the pore helix and the second turn of the pore helix (Fig 6c), and two inter-subunit hydrogen bonds, Tyr4888-Arg4913 and Tyr4888-Asp4917 between the pore helix and S6. These three interactions may stabilize the luminal mouth of the channel and hold S6 through its conformational transitions.

## Discussion

The goal of this work was to understand the mechanism of Ca^2+^-induced inactivation of RyR1 from a structural point of view. Under our conditions of 2 mM free Ca^2+^, we discerned two classes within each of the two datasets, with their ion gate either in an open or a closed conformation on account of direct pore analysis. 2 mM free Ca^2+^ concentration is near the Ca^2+^ IC_50_ found in the [^3^H] ryanodine binding assays in the presence of ATP analogue, a concentration at which Ca^2+^-activated and Ca^2+^-inactivated RyR1 conformations should coexist. Accordingly, we propose that they correspond with the open-pore and closed-pore conformations observed by cryo-EM, respectively.

Earlier 3D structures of RyR in the open state required synthetic activators such as caffeine, which by itself suffices to activate the channel (Xu *et al*, 2018), or PCB95 (Bai et al., 2016, Samsó et al., 2009). Our cryo-EM maps of RyR1 and lipid were obtained in the absence of any non-physiological activator, and in a nanodisc environment. The two independently structures of RyR1-ACP/Ca^2+^ open (A and B) obtained in the presence of Ca^2+^ and ACP provide a more accurate account of the native open state and overall, validate the previous structures obtained using artificial agonists.

Activation and inactivation appear to proceed under an integrated mechanism, where inactivation depends on prior activation. Both RyR1 open and inactivated conformations showed Ca^2+^ bound at the high affinity site in the CD/CTD interface, which joined these and their proximal domains (EF, U-motif, and S6C’) into a more rigid block. In activation, CD and CTD rotate towards each other closing around the Ca^2+^ ion; the rotation of the CTD pulls S6C’ outwards, opening the pore. In inactivation, the CD and connected central rigid block rotate further into the arc initiated by the CD alone, which now pushes the S6 helices towards the central axis. The movement is similar to pushing down the levers of a winged corkscrew while pushing towards the central rod (see schematics in Fig 2d), closing the channel in a distinctive conformation different from the EGTA-closed resting state. This state, with a closed pore and Ca^2+^ bound to the activation site, can no longer be activated by Ca^2+^, which is the hallmark of an inactivated state.

Comparison of the open, closed and inactivated conformations uncovers other novel features. A strong electron density connected to ACP under the high Ca^2+^ conditions, that we attribute to hydrated Ca^2+^, progressively supports more interactions with the CTD and S6, increasing cohesiveness of the central block (in the order inactivated > activated > closed; Fig 4), which could explain the more robust Ca^2+^-induced inactivation reported in the presence of ATP (Sitsapesan & Williams, 2000). Considering that Mg^2+^ at its physiological concentration of ~5 mM probably interacts with RyR1-bound ATP, this likely constitutes a secondary binding site for divalent cations. At least in the resolved regions of the RyR1 structure, there were no other obvious densities that could account for a bound Ca^2+^ ion, or clusters of negatively charged residues that could support Ca^2+^-mediated conformational changes.

The EF hand domain did not have Ca^2+^ bound in the high Ca^2+^ datasets in keeping with the low affinity of Ca^2+^ to this site (Xiong et al., 1998). However, this protruding domain, with its sequence between the CD and the U-motif, appears to play a distinctive role in inactivation which derives from its further anticlockwise and out-of-plane rotations with respect to the open state (Fig 3). This conformation brings the EF hand domain in contact with the cytoplasmic loop between the S2 and S3 transmembrane helices of the neighboring subunit, forming two inter-subunit electrostatic interactions (Fig 3). One possible scenario is that the energy landscape of the open channel allows for overshoot of the opening motion, allowing interaction of the EF hand and S2-S3 loop domains by such salt bridges. These appear to stabilize the inactivated conformation by providing an extra linkage between subunits, and between the cytoplasmic assembly and TMD.

In the TMD, two lipids in a crevice between S3/S4 and S5/S6 of adjacent subunits were resolved in the inactivated state. The hydrophobic nature of this crevice suggests that lipids may help to stabilize the TMD. In the open state the hydrophobic crevice is narrower, and Phe4808 of S4 adopts a different orientation that would clash with lipid-2, suggesting rearrangement in the open state. A similar lipid exclusion of bound lipid in the open channel was also reported for the TRPV3 channel (Singh et al., 2018). Departure from α-helix geometry was also present in segments of S4 (near-3_10_ helix and π helix), and S5, S6 (π helix), with obvious correlation with the closed, open and inactivated conformations in S4. The dynamism of the transmembrane helices through the gating transitions appears to be supported by anchoring of Phe motifs in S4, S5 and S6 to lipids in the membrane. In addition, inter-subunit interactions including disulfide bridges link the 4860-4878 luminal loop, the pore helix and S6N’ around the luminal mouth of the channel where the four protomers converge.

Functional studies employing single channels embedded in lipid bilayers show rapid channel inactivation following an RyR1 Ca^2+^ release event. Therefore, after RyR1 opening, a refractory period is needed to relieve inactivation and recover the ability to activate again (Laver & Lamb, 1998, Ríos et al., 2008, Schiefer *et al*, 1995, Sitsapesan & Williams, 2000). The 3D reconstructions reported here provide a structural basis for this refractoriness: the Ca^2+^-inactivated state is in a distinctive locked closed conformation while the high affinity Ca^2+^ binding site is still occupied. This renders this conformation unable to be activated by Ca^2+^, as long as Ca^2+^ occupies the high affinity site. In this sense, Ca^2+^permeation through the RyR1 would provide negative feedback. Ca^2+^-dependent inactivation is often observed in Ca^2+^ permeation pathways, probably to limit cytosolic Ca^2+^ overload that could be detrimental and life-threatening (Dick *et al*, 2016, Gomez et al., 2016). Here we provide a structural basis to understand the transitions from closed, to open and then to the inactivated state of the RyR1 at high resolution, and together with previous investigations (Gomez et al., 2016, Gomez & Yamaguchi, 2014), we propose a structural mechanism for how naturally occurring mutations disturb RyR1 inactivation producing Ca^2+^ dysregulation and muscle disease.

## Materials and Methods

### Reagents

All chemicals were purchased from ThermoFisher or Sigma-Aldrich except where indicated.

### [^3^H]Ryanodine binding

RyR1 activity was estimated by measuring the extent of bound ^3^[H]ryanodine in microsomes isolated from rabbit skeletal muscle when incubated with free Ca^2+^ alone (10 μM to 2 mM range), or in the presence of 2 mM ATP or 2 mM AMP-PCP (ACP). Concentrations of total Ca^2+^ added to the reaction mixture were estimated in Maxchelator (https://somapp.ucdmc.ucdavis.edu/pharmacology/bers/maxchelator). Preincubated membrane vesicles (~40 μg) were allowed to bind 5 nM [^3^H]ryanodine (PerkinElmer) in a buffer containing 50 mM MOPS (pH 7.4), 0.15 M KCl, 0.3 mM EGTA, protease inhibitors, and 2 mM DTT for 3 h at 37°C. Sample aliquots were diluted seven-fold with an ice-cold wash buffer (0.1 M KCl) before placing onto Whatman GF/B filter papers in a vacuum-operated filtration apparatus. The remaining radioactivity in the filter papers after washing three times with the wash buffer was measured by liquid scintillation counting. Non-specific ryanodine binding was estimated in the presence of 250 μM unlabeled ryanodine (Calbiochem) and subtracted from the total binding. Data represent the mean specific [^3^H] ryanodine binding from four independent experiments.

### Purification of RyR1 from rabbit skeletal muscle and reconstitution into nanodiscs

Microsomes were purified from rabbit back and hind leg muscles through differential centrifugation as previously described (Hu *et al*, 2021, Samsó et al., 2009). 100 mg frozen membranes were thawed and solubilized in buffer-A containing 20 mM MOPS pH 7.4, 1 M NaCl, 9.2% (w/v) CHAPS, 2.3% (w/v) Phosphatdylcholine (PC; Sigma), 2 mM DTT and protease inhibitor cocktail for 15 mins at 4° C. The solubilized membranes were centrifuged at 100,000 x g for 60 min and the pellet was discarded. Supernatant was layered onto 10-20% (w/v) discontinuous sucrose gradients, prepared in buffer-A containing 0.5% CHAPS and 0.125% PC (buffer-B). The layered sucrose gradient tubes were ultracentrifuged at 120,000 x g for ~20 h at 4° C to allow RyR1 separation. Fractions containing >95% pure RyR1 were pooled and further purified with a HiTrap Heparin HP Agarose column (GE Healthcare), after a five-fold dilution and filtration step. RyR1 was eluted with buffer B containing 0.9 M NaCl after washing with 20 column volumes of buffer B with 200 mM NaCl. Peak fractions were flash-frozen and stored at −80° C until reconstitution into nanodiscs and cryo-EM. 1.5-2 mg of RyR1 was purified from 100 mg of SR membrane vesicles. RyR1 purity was estimated with 12.5% SDS-PAGE and negative staining with 0.75% uranyl formate. Protein concentration in purified microsomes and RyR1 fractions was measured with Quick Start Bradford Protein Assay (Bio-Rad). The plasmid encoding for MSP1E3D1, pMSP1E3D1, was purchased from Addgene and recombinant MSP1E3D1 was purified in *E. coli* using the manufacturer’s instructions. RyR1-nanodiscs were obtained by mixing purified RyR1, MSP1E3D1 and POPC (Avanti polar lipids) at a 1:2:50 molar ratio. The mixture was incubated for 1 h 30 min at 4° C before an overnight dialysis in a CHAPS-free buffer (20 mM MOPS pH 7.4, 635 mM KCl, 2 mM DTT), which contained either 1 mM EGTA + 1mM EDTA for the control “RyR1-ACP/EGTA” dataset or 3.7 mM CaCl2 for the RyR1-ACP/ Ca^2+^ dataset. The dialyzed RyR1-nanodisc preparations were incubated with ACP-PCP for 30 min prior to plunge freezing, at concentrations of 5 mM ACP (control RyR1-ACP/EGTA dataset) or 2 mM ACP (RyR1-ACP/Ca^2+^datasets). Free Ca^2+^ was calculated with Maxchelator. Integrity of the nanodisc-embedded channels was examined by negative staining.

### Cryo-EM grid preparation and data acquisition

Cryo-EM grids were cleaned with a customized protocol (Passmore & Russo, 2016) prior to glow discharge. Aliquots of 1.25 – 1.5 μl RyR1-Nanodisc were applied onto each side of glow-discharged UltraAufoil −1.2/1.3 holey-gold (Quantifoil, Germany) with 300 TEM mesh. The grids were blotted for 1-1.5 s with an ashless Whatman^®^ Grade 540 filter paper in a Vitrobot Mark IV (ThermoFisher Scientific) and rapidly plunged into liquid ethane. Grid quality and RyR1 sample distribution was assessed on a Tecnai F20 (ThermoFisher Scientific) electron microscope. Data acquisition was carried out in a Titan Krios transmission electron microscope (ThermoFisher Scientific) operated at 300 kV and counting mode, with a K2 or K3 detector (Gatan), in two separate sessions for ACP/Ca^2+^_A_ and ACP/Ca^2+^_B_ datasets respectively. A Gatan Quantum Energy Filter (GIF) with a slit width of 20 eV was employed. The ACP/EGTA was collected on a K2 detector and a 20eV GIF in a separate session. Datasets were collected in automated mode with the program Latitude (Gatan) with cumulative electron dose of 70 e^−^/Å^2^ applied over 50-60 frames. Image acquisition parameters for RyR1-ACP/EGTA, RyR1-ACP/Ca^2+^_A_ and RyR2-ACP/Ca^2+^_B_ datasets are summarized in Table S1.

### Single-Particle Image Processing

Gain-reference normalization, movie-frame alignment, dose-weighting, motion correction of the collected movie stacks was carried out with Motioncor2 (Zheng *et al*, 2017). Contrast transfer function parameters were estimated from non-dose weighted motion corrected images using Gctf (Zhang, 2016). All subsequent image processing operations were carried out using dose-weighted, motion-corrected micrographs in RELION 3.0 (Scheres, 2012). The micrographs were low-pass filtered to 20 Å before automated particle picking. 2D class average templates for autopicking were generated by reference-free 2D classification of 1000 manually picked particles. Autopicked particles with ethane and hexagonal ice-contaminated areas were removed by visual inspection. Particle image sub-stacks required for the focused reconstructions residues 3668-5037, encompassing the CD, U-motif, TMD and CTD domains, were generated using a signal subtraction procedure employed in relion_project module (Bai *et al*, 2015). Particle image stacks of quarter sub-volumes of RyR1 corresponding to a single subunit were generated by particle subtraction following a symmetry expansion step with relion_particle_symmetry_expand tool (Bai et al., 2015, Scheres, 2016). Composite tetrameric maps were generated from the symmetry expanded monomeric maps with chimera (Pettersen *et al*, 2004) vop maximum tool. B-factor applied to the reconstructed maps was estimated with relion_postprocess. Unfiltered half maps obtained from the final 3D-refinement step in RELION 3.0 were further density modified with Phenix.Resolve (Terwilliger et al., 2020). The reported resolutions of the cryo-EM maps are based on FSC 0.143 criterion (Scheres & Chen, 2012). Local resolution was estimated with ResMap (Kucukelbir *et al*, 2014). Pixel size calibration of post-processed maps were carried out using real space correlation metric of UCSF Chimera using a published RyR1 cryo-EM map (des Georges et al., 2016). Pixel size maxima of 1.07, 1.105 and 1.07 Å were obtained in RyR1-ACP/EGTA, RyR1-ACP/Ca^2+^_A_, RyR1-ACP/Ca^2+^_B_ respectively. Image processing schemes of the RyR1-ACP/Ca^2+^_A_, RyR1-ACP/Ca^2+^_B_, RyR1-ACP/EGTA datasets are summarized in Figs S1, S3 and S4, respectively.

### Model Building and Structure refinement

The cryo-EM based atomic models of RyR1 (PDB ID: 5tb3 for RyR1-ACP/EGTA, RyR1-ACP/Ca^2+^inactivated and 5ta3 for RyR1-ACP/Ca^2+^ open) were taken as the initial models for model building. The best resolved symmetry expanded cryo-EM map for a single subunit in inactivated or open conformation was docked within a RyR1 monomer model with Chimera Fit in map tool. Local density fit of the RyR1 sequence was improved over an iterative process of amino acid fitting in coot (Emsley *et al*, 2010) alternated with real space refinement in PHENIX (Afonine *et al*, 2018). Four copies of the monomers were docked to the whole RyR1 reconstructions. Real space refinement of the tetrameric models was carried out with secondary structure and Ramachandran restraints. Further manual fitting of the CD, TMD and CTD (3668-5037) of RyR1 was carried out in Coot. Comprehensive model validation was carried out with PHENIX and PDB validation server at https://validate-rcsb-2.wwpdb.org/ and are summarized in Table S2. Molecular lipophilicity potential surfaces were drawn in chimeraX (Ghose *et al*, 1998, Laguerre *et al*, 1997, Pettersen *et al*, 2021). Figures were generated with PYMOL (Schrodinger, 2015) and chimera programs (Pettersen et al., 2004, Pettersen et al., 2021).

### Pore radius and Helical geometry measurements

Pore radii were measured for the refined atomic model coordinates of the RyR1 pore region (residues 4821-5037) with the HOLE program (Smart *et al*, 1993). Dot-surfaces representing the channel ion permeation pathway were generated with HOLE implemented in Coot, which were reformatted to enable visualization in UCSF chimera. Residues per turn of S4 and S6 transmembrane helices of RyR1 were calculated with HELANAL (Bansal *et al*, 2000).

### Flexion angle measurement

The cytoplasmic shell flexion angles of RyR1 in different conformations were estimated using (Steele & Samsó, 2019) with minor changes. The angle was calculated between a diagonal running from the N-terminal domain (residue 348) to the P1 domain (residue 984) from the same subunit, and the horizontal plane.

### Data availability

Tetrameric and focused cryo-EM maps of RyR1-ACP/EGTA, RyR1-ACP/Ca^2+^inactivated and open of dataset A and B have been deposited in the Electron Microscopy Databank (EMDB) with accession codes 22616, 22597 (RyR1-ACP/EGTA), X, and Y, respectively. Atomic models generated from cryo-EM maps of RyR1-ACP/EGTA, RyR1-ACP/Ca^2+^_A_ inactivated and RyR1-ACP/Ca^2+^_A_ open have been deposited in the RCSB PDB database with accession codes 7K0T, Y, and Z, respectively.

## Appendix

Supplementary material for this article is available in a separate file.

## Acknowledgments

Cryo-grid preparation and screening were carried out at the Cryo-EM Unit at Virginia Commonwealth University (VCU) supported by the VCU School of Medicine and M.S.’s funds. Cryo-EM data collection was carried out at the Frederick National Laboratory for Cancer Research supported by contract HSSN261200800001E, at the Molecular Electron Microscopy Core Facility at the University of Virginia (UVA) (supported by NIH U24 GM116790). We thank Drs. Thomas Edwards, Ulrich Baxa and Adam Wier for cryo-EM data collection at the Frederick National Laboratory. Supported by NIH R01 AR068431 and the Muscular Dystrophy Association (MDA 352845) (to M.S.).

## Author contributions

A.R.N. performed protein purification, cryo-grid preparation, image processing, model building and radioligand binding; M.S. conceived, designed and supervised all experiments, M.S. and A.R.N. interpreted the data and wrote the manuscript.

