## Supplementary Material for "Ca^2+^-inactivation of the mammalian ryanodine receptor type 1 in a lipidic environment revealed by cryo-EM"

**Table of Contents**

Figures S1 to S9
Tables S1 to S2
Movies S1 to S2

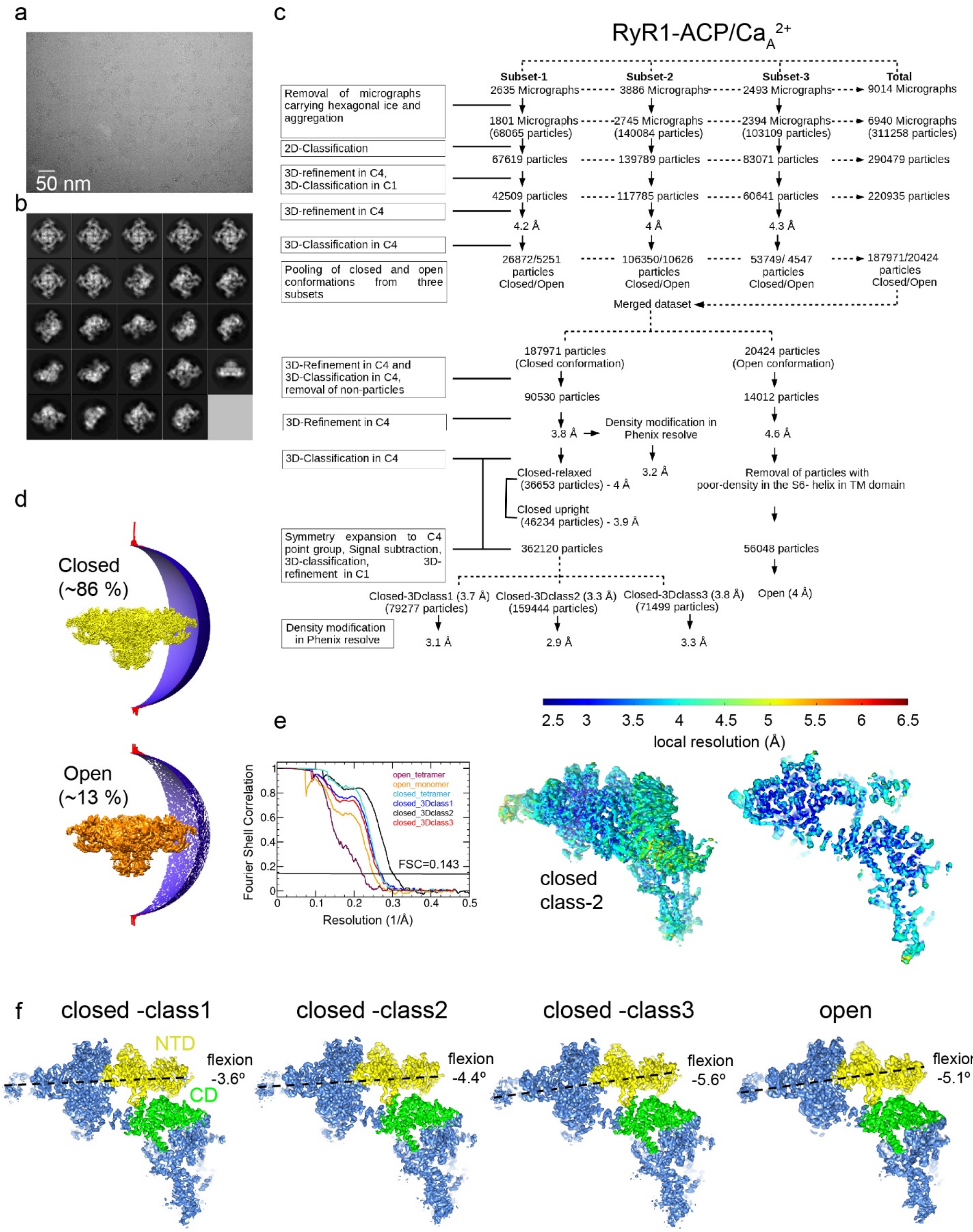
a
50 nm
b
c
RyR1-ACP/CaA2+
Subset-1
Subset-2
Subset-3
Total
Removal of micrographs carrying hexagonal ice and aggregation
2D-Classification
3D-refinement in C4, 3D-Classification in C1
3D-refinement in C4
3D-Classification in C4
Pooling of closed and open conformations from three subsets
2635 Micrographs
3886 Micrographs
2493 Micrographs
9014 Micrographs
1801 Micrographs (68065 particles)
2745 Micrographs (140084 particles)
2394 Micrographs (103109 particles)
6940 Micrographs (311258 particles)
67619 particles
139789 particles
83071 particles
290479 particles
42509 particles
117785 particles
60641 particles
220935 particles
4.2 Å
4 Å
4.3 Å
26872/5251 particles Closed/Open
106350/10626 particles Closed/Open
53749/ 4547 particles Closed/Open
187971/20424 particles Closed/Open
Merged dataset
3D-Refinement in C4 and 3D-Classification in C4, removal of non-particles
3D-Refinement in C4
3D-Classification in C4
Symmetry expansion to C4 point group, Signal subtraction, 3D-classification, 3D-refinement in C1
Density modification in Phenix resolve
187971 particles (Closed conformation)
20424 particles (Open conformation)
90530 particles
14012 particles
3.8 Å
Density modification in Phenix resolve
4.6 Å
3.2 Å
Closed-relaxed (36653 particles) - 4 Å
Closed upright (46234 particles) - 3.9 Å
Removal of particles with poor-density in the S6- helix in TM domain
362120 particles
56048 particles
Closed-3Dclass1 (3.7 Å) (79277 particles)
Closed-3Dclass2 (3.3 Å) (159444 particles)
Closed-3Dclass3 (3.8 Å) (71499 particles)
Open (4 Å)
3.1 Å
2.9 Å
3.3 Å
d
Closed (~86 %)
Open (~13 %)
e
open_tetramer
open_monomer
closed_tetramer
closed_3Dclass1
closed_3Dclass2
closed_3Dclass3
FSC=0.143
Fourier Shell Correlation
Resolution (1/Å)
2.5 3 3.5 4 4.5 5 5.5 6 6.5
local resolution (Å)
closed class-2
f
closed -class1
closed -class2
closed -class3
open
NTD
CD
flexion -3.6º
flexion -4.4º
flexion -5.6º
flexion -5.1º

**Figure S1. Image Processing Scheme for the RyR1-ACP/$Ca^{2+}{}_A$ dataset**

**a,** Representative micrograph collected on a Titan Krios at 81,000x magnification with a K3 camera in counting mode. **b,** Representative 2D class averages obtained by reference-free 2D classification in RELION. **c,** Image processing workflow with tetrameric and single subunit-masked RyR1 particles in RELION. Overall resolution values obtained for the inactivated conformations after density modification in Phenix resolve. **d,** Euler angle distribution of the particles contributing to the RyR1-ACP/$Ca^{2+}$ inactivated (3.8 Å resolution) and RyR1-ACP/$Ca^{2+}$ open (4.6 Å resolution) maps. **e,** Left, Gold standard Fourier shell correlation prior to density modification. Right, Local resolution of the symmetry-expanded maps calculated with ResMap. **f,** Main 3D classes obtained from the focused 3D classification after symmetry expansion with NTD and CD domains highlighted in yellow and green, respectively, with the pore axis on the right. Classification yielded one open conformation and three closed conformations that differ in the flexion angle of the cytoplasmic domain.

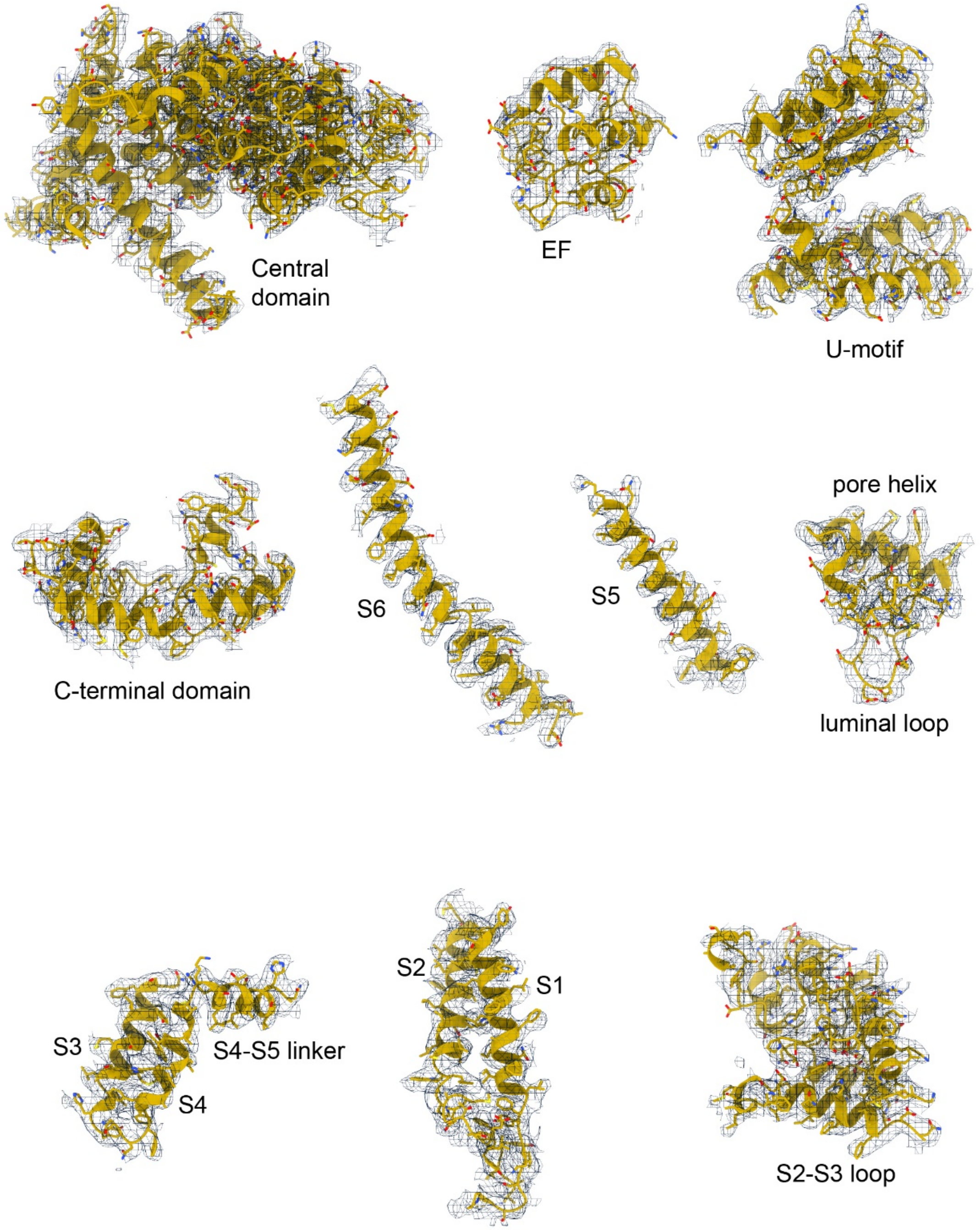


**Figure S2. Correlation of RyR1-ACP/$Ca^{2+}{}_A$ inactivated model with the cryo-EM density**

Model quality for RyR1-ACP/$Ca^{2+}{}_A$ inactivated in the consensus cryo-EM map; domains in the central and transmembrane regions are shown.

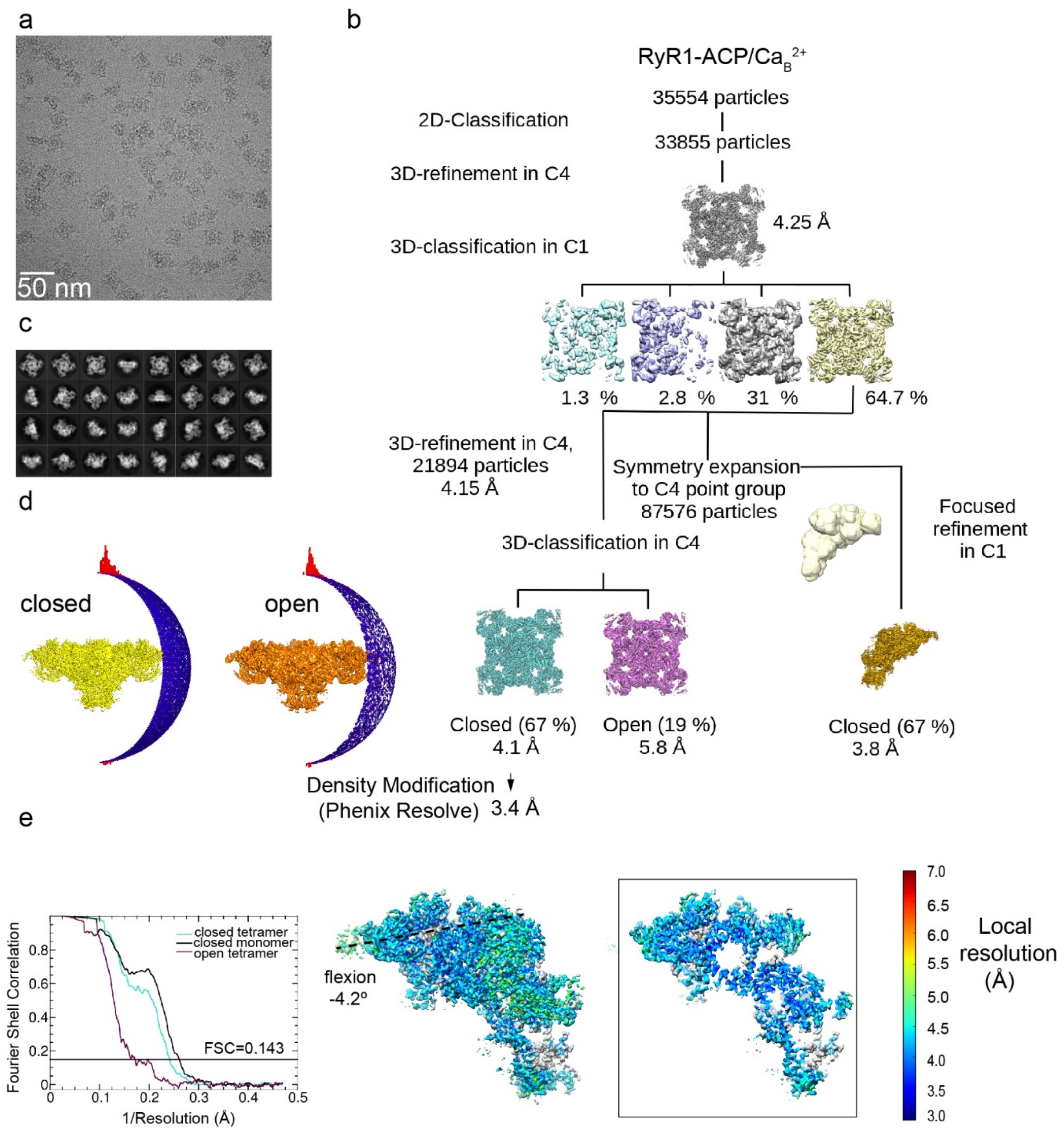


**Figure S3. Image processing scheme for the RyR1-ACP/$Ca^{2+}_B$ dataset**

**a,** Representative micrograph collected on a Titan Krios at 130,000x magnification with a K2 camera in super resolution mode. **b,** Representative 2D class averages obtained by reference-free 2D classification in RELION. Overall resolution values obtained for inactivated 3D-class after a density modification in Phenix resolve. **c,** Workflow of image processing with tetramer and single subunit-masked particles carried out in RELION. **d,** Euler angle distribution of the particles contributing to the RyR1-ACP/$Ca^{2+}_B$ inactivated and RyR1-ACP/$Ca^{2+}_B$ open maps at 4.1 Å and 5.8 Å resolution, respectively. **e,** Left, Gold standard Fourier shell correlation prior to density modification. Right, Local resolution of the symmetry-expanded map calculated with ResMap with flexion angle of the cytoplasmic domain indicate

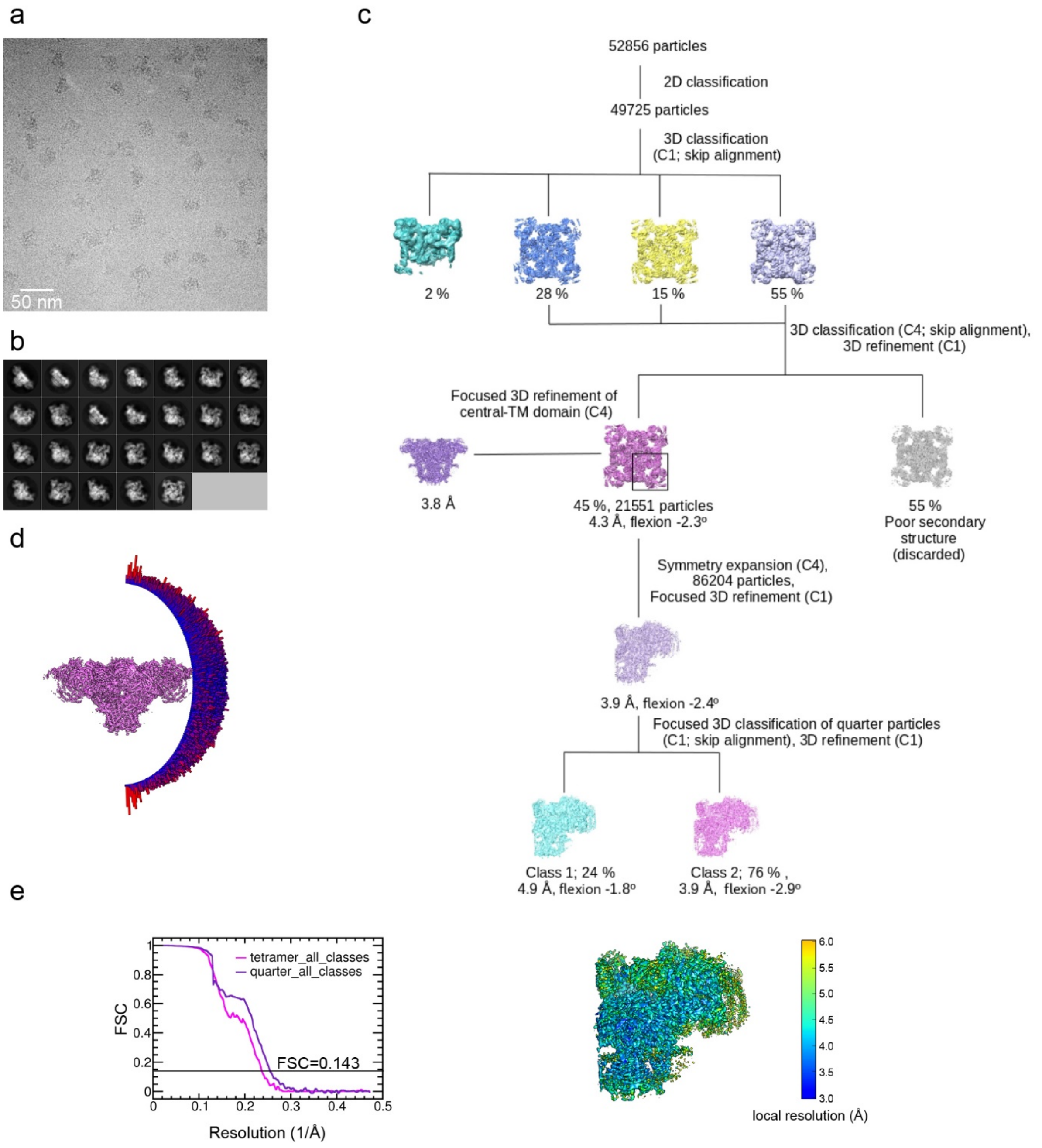


**Figure S4. Image processing scheme for the RyR1-ACP/EGTA dataset**

**a,** Representative micrograph collected on a Titan Krios at 130,000x magnification with a K2 camera in super-resolution mode. **b,** Representative 2D class averages obtained by reference-free 2D classification in RELION. **c,** Image processing workflow including 3D classification focused on RyR1's quarter carried out in RELION. **d,** Euler angle distribution of the particles contributing to the 4.3 Å resolution 3D map. **e,** Left, Gold standard Fourier shell correlation. Right, Local resolution of the symmetry-expanded map calculated with ResMap.

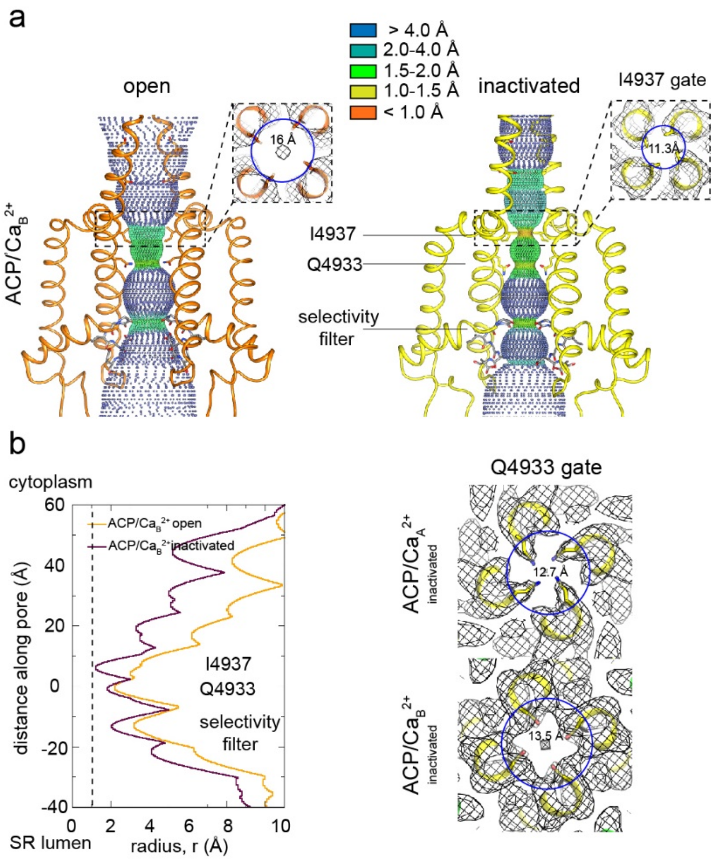


**Figure S5. Pore conformation in the RyR1-ACP/Ca$^{2+}_B$ dataset**

**a,** Dotted surfaces of RyR1 ion-permeation pathway in open and inactivated conformations. Inset, Fourfold cytoplasmic views of Ile4937 constriction and corresponding pore diameter (measured at Cα backbone) with respective cryo-EM maps (mesh). **b,** Left, Pore profiles of inactivated and open conformations calculated with HOLE. Radius corresponding to a dehydrated $Ca^{2+}$ ion is shown as a dotted line. Right, fourfold cytoplasmic views of the Gln4933 constriction in RyR1-ACP/Ca$^{2+}_A$ and RyR1-ACP/Ca$^{2+}_B$ pores with their respective cryo-EM maps.

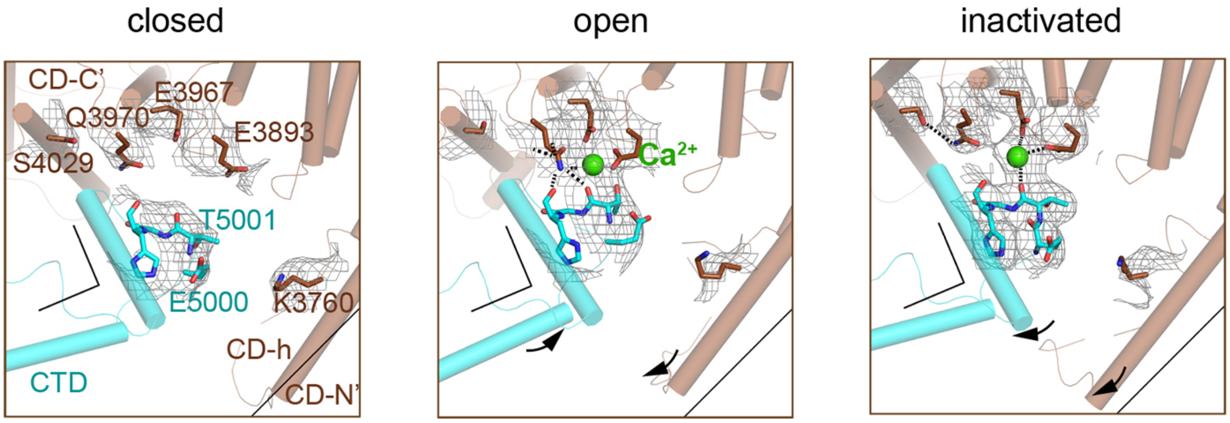


**Figure S6. Reorganization of the high-affinity $Ca^{2+}$ binding site at the CD/CTD interface under different conditions**

The high-affinity $Ca^{2+}$ binding site with electron density around the $Ca^{2+}$ site contoured at 4σ. Contacts within 2.8 Å from $Ca^{2+}$, as well as additional contact Gln3970-Ser4029 within 3.6 Å are represented by dashed lines. Channel axis is on the left. During the transition from open to inactivated conformations, the CD/CTD block tilts around the $Ca^{2+}$ binding site such that the protruding fourth helix of the CD (CD-h) and connected CTD tilt inward, while the CD-C' tilts upward and away from the SR membrane -with Gln3970 separating from $Ca^{2+}$ by ~6 Å. Arrows and stationary reference lines illustrate the conformational changes undergone with respect to the panel on the left. The region represented relative to the channel is highlighted with a square in Fig 2, panel c.

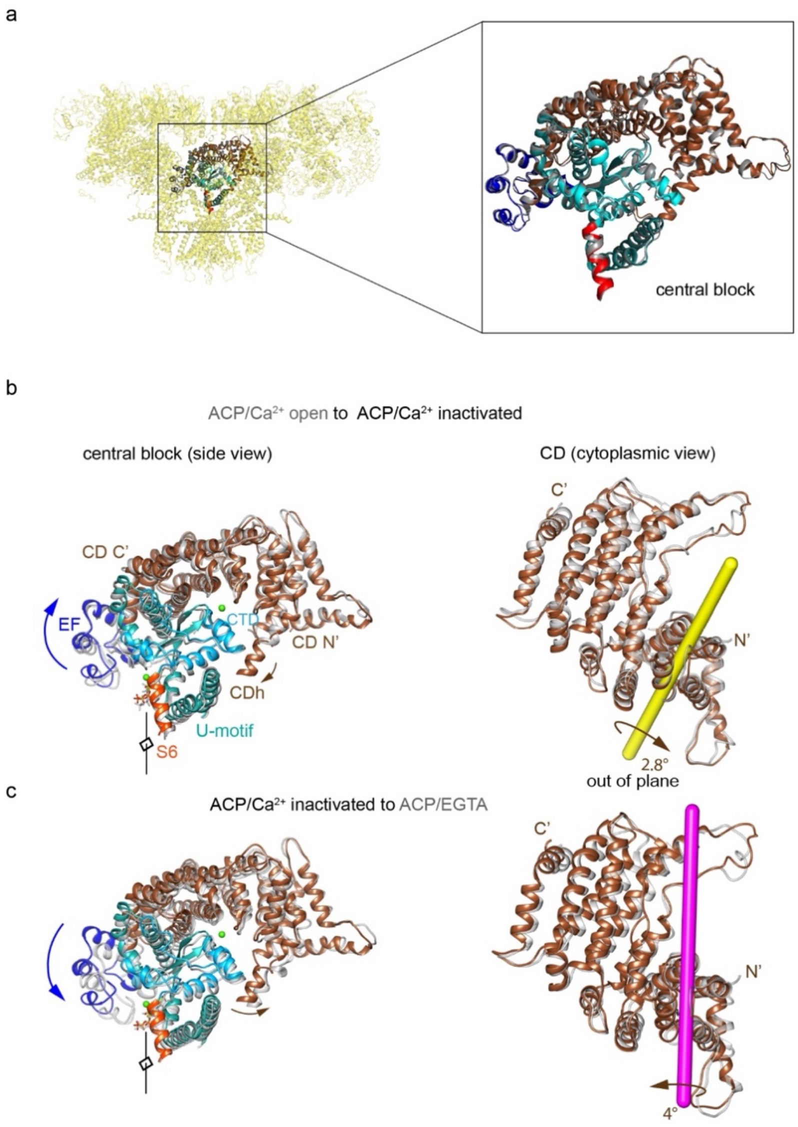


**Figure S7. Rotation axes of the CD of RyR1 among the different conformational transitions**
**a,** Reproducibility of the conformation of the "central block" of RyR1-ACP/$Ca^{2+}$-inactivated from two independent datasets (superimposed), with an RMSD of 0.94 Å over the 672 Cα atoms. We define the central block as residues 3668-4251 (CD and associated domains EF and U-motif) together with residues 4945-5037 (CTD and associated S6C'). **b,** Transition from ACP/$Ca^{2+}$ open (gray) to ACP/$Ca^{2+}$ inactivated (colored) involves an out-of-plane rotation of the CD around the yellow-colored axis. This results in inward tilt in CD-N' (brown arrows) and upward shift/tilt of CD-C' and connected EF hand domain (blue arrows). Two orthogonal views are shown, and the fourfold axis on the left panels, in front of the structure, is indicated. **c,** Transition from ACP/$Ca^{2+}$ inactivated (colored) to ACP/EGTA closed (gray) involves out-of-plane rotation of the CD of RyR1 around the magenta-colored axis.

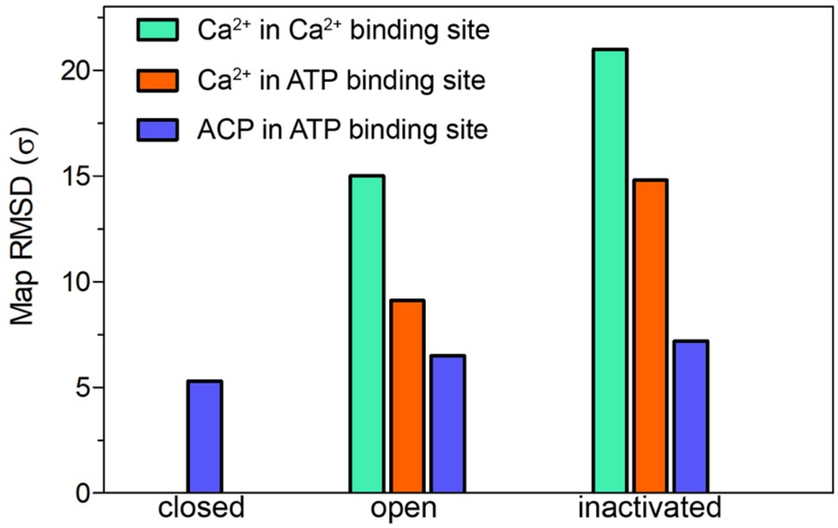


**Figure S8. Map significance of the ACP and putative $Ca^{2+}$ densities under different conditions**

Root Mean Square Deviation (RMSD σ) values of the densities attributed to $Ca^{2+}$ and ACP in the cryo-EM maps of RyR1-ACP/EGTA closed, RyR1-ACP/$Ca^{2+}_A$ open and RyR1-ACP/$Ca^{2+}_A$ inactivated.

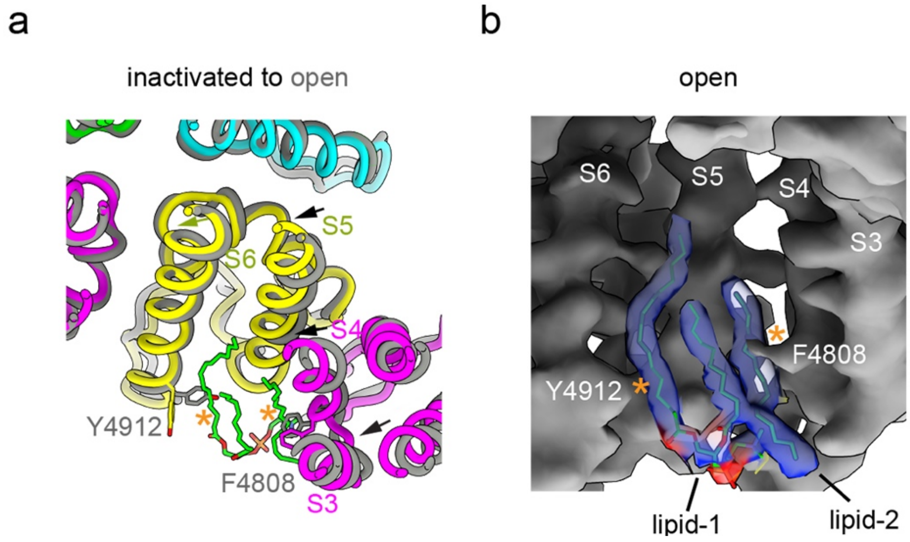


**Figure S9**. **Lipids as bound to the TMD crevice in the inactivated conformation would cause steric clash in the open conformation**

**a,** Cytoplasmic fourfold view of part of the TMD showing movement of the transmembrane helices S3, S4 and S5 (arrows) into the lipid pocket while transitioning from ACP/$Ca^{2+}{}_A$ open to inactivated conformations. No ordered lipids were observed in the open channel. **b,** Isosurface of the lipophilic pocket in RyR1-ACP/$Ca^{2+}{}_A$ open with overlaid lipids extracted from the RyR1-ACP/$Ca^{2+}{}_A$ inactivated conformation, showing steric hindrance of lipid-1and lipid-2 with S6 and S4, respectively. See also Fig 6a.

### Table S1. Summary of cryo-EM data collection and image processing parameters

| Dataset | RyR1 ACP/EGTA closed | RyR1 ACP/$Ca^{2+}_A$ inactivated | RyR1 ACP/$Ca^{2+}_A$ inactivated class1 | RyR1 ACP/$Ca^{2+}_A$ inactivated class2 | RyR1 ACP/$Ca^{2+}_A$ inactivated class3 | RyR1 ACP/$Ca^{2+}_A$ inactivated CDTM | RyR1 ACP/$Ca^{2+}_A$ open | RyR1 ACP/$Ca^{2+}_A$ open CDTM | RyR1 ACP/$Ca^{2+}_B$ inactivated | RyR1 ACP/$Ca^{2+}_B$ open |
|---|---|---|---|---|---|---|---|---|---|---|
| **Data Acquisition** | | | | | | | | | | |
| Microscope/Detector | Krios/K2 | Krios/K3 | | | | | | | Krios/K2 | |
| Voltage (kV) | 300 | 300 | | | | | | | 300 | |
| Magnification | 130000 | 81000 | | | | | | | 130000 | |
| Defocus range (μm) | -1.2 to -2.2 | -1.25 to -2.5 | | | | | | | -1.25 to -2.5 | |
| Pixel Size (Å) (calibrated) | 1.06 (1.07) | 1.08 (1.105) | | | | | | | 1.06 (1.07) | |
| Total electron dose (e/$Å^2$) | 70 | 70 | | | | | | | 70 | |
| Exposure time (sec) | 12 | 4.4 | | | | | | | 14 | |
| Number of frames | 60 | 50 | | | | | | | 50 | |
| Total number of Micrographs | 1,959 | 10002 | | | | | | | 1346 | |
| **Image Processing** | | | | | | | | | | |
| Total number of Particles selected | 52,856 | 311258 | | | | | | | 35554 | |
| Final number of Particles | 21,551 | 90530 | 79277 | 159444 | 71499 | 90530 | 14012 | 14012 | 14669 | 4160 |
| Reconstruction symmetry | C4 | C4 | C1 | C1 | C1 | C4 | C4 | C4 | C4 | C4 |
| Map Resolution, FSC (0.143) (Å) (symmetry expanded map) | 4.1(3.9) | 3.8 (3.5) | 3.7 | 3.3 | 3.8 | 3.5 | 4.6 (4.0) | 4.4 | 4.1 (3.8) | 5.8 |
| Map sharpening B-factor ($Å^2$) (symmetry expanded map) | -132 | -150 (-144) | -125 | -102 | -123 | -142 | -168 (-135) | -144 | -111(-89) | -148 |
| EMDB ID | 22616,22597 | | | | | | | | | |

### Table S2. Summary of model refinement and validation statistics

| Dataset | RyR1 ACP-EGTA | RyR1 ACP/$Ca^{2+}$A inactivated | RyR1 ACP/$Ca^{2+}$ A inactivated class1 | RyR1 ACP/$Ca^{2+}$ A inactivated class2 | RyR1 ACP/$Ca^{2+}$ A inactivated class3 | RyR1 ACP/$Ca^{2+}$ A inactivated CDTM | RyR1 ACP/$Ca^{2+}$ A open | RyR1 ACP/$Ca^{2+}$ B inactivated |
|---|---|---|---|---|---|---|---|---|
| RMS Deviation (Bonds) | 0.004 | 0.002 | 0.002 | 0.002 | 0.002 | 0.004 | 0.012 | 0.007 |
| RMS Deviation (Angle) | 0.633 | 0.583 | 0.594 | 0.581 | 0.584 | 0.651 | 0.605 | 1.006 |
| Ramachandran Plot statistics (%) | | | | | | | | |
| Preferred | 92.41 | 93.14 | 93.57 | 93.55 | 93.6 | 93.09 | 92.69 | 86.9 |
| Allowed | 7.43 | 6.86 | 6.8 | 6.45 | 6.4 | 6.91 | 7.29 | 12.98 |
| Outliers | 0.16 | 0.01 | 0 | 0 | 0 | 0 | 0.02 | 0.12 |
| Clash-score | 7.39 | 7.1 | 6.4 | 6.3 | 6.4 | 5.95 | 9.83 | 4.8 |
| MolProbity Score | 1.82 | 1.83 | 1.8 | 1.77 | 1.78 | 1.77 | 1.98 | 1.87 |
| PDB ID | 7K0T | | | | | | | |

**Movie S1. $Ca^{2+}$-induced transitions in activation and inactivation**

In activation, $Ca^{2+}$ binding joins CD and CTD-S6C' region, separating the S6 four-helix bundle and opening the channel. High $Ca^{2+}$-induced inactivation promotes out-of-plane rotation of the CD-CTD-S6C' block. This pushes in the S6 helices, closing the channel. Straightening of S6 upon closure results in transition from a wide to a narrow π-helix around the S6N' region (4920-4928). Only two protomers in diagonal are shown.

**Movie S2. Gating-induced conformational changes in nanodisc**

Conformational change in upper and lower nanodisc belts while morphing from RyR1-ACP/$Ca^{2+}{}_{A}$ open to inactivated conformations; the view translates from the upper-belt (positioned at the S4-S5 linker level) to the lower-belt towards the luminal side. The nanodisc, low-pass filtered to 7 Å resolution, can be seen expanding when transitioning to the open state.
